# Cell-type specific population codes link inferior temporal cortex to object recognition behavior

**DOI:** 10.1101/2023.08.01.551579

**Authors:** Sabine Muzellec, Sachi Sanghavi, Kohitij Kar

**Affiliations:** Department of Biology, Faculty of Science, Centre for Vision Research, and Centre for Integrative and Applied Neuroscience, York University, Toronto, Ontario, Canada; Department of Computer Sciences, University of Wisconsin-Madison, USA

**Author notes:** denotes equal contribution (Sabine Muzellec, Sachi Sanghavi). Correspondence should be addressed to Kohitij Kar, Sabine Muzellec.

**Keywords:** Inferior temporal cortex, cell types, population decoding, deep neural networks, object recognition

## Abstract

A spatially distributed population of neurons in the macaque inferior temporal (IT) cortex supports object recognition behavior, but the cell-type specificity of the population in forming "behaviorally sufficient" object decodes remains poorly understood. To address this, we recorded neural signals from the macaque IT cortex and compared the object identity information and the alignment of decoding strategies derived from putative inhibitory (Inh) and excitatory (Exc) neurons to the monkeys’ behavior. We observed that while Inh neurons represented significant category information, decoding strategies based on Exc neural population activity outperformed those from Inh neurons in overall accuracy and their image-level match to the monkeys’ behavioral reports. Interestingly, both Exc and Inh responses explained a fraction of unique variance of the monkeys’ behavior, demonstrating a distinct role of the two cell types in generating object identity solutions for a downstream readout. We observed that current artificial neural network (ANN) models of primate ventral stream, designed with AI goals of performance optimization on image categorization, better predict Exc neurons (and their contribution to object recognition behavior) than Inh neurons. Beyond, refining the linking propositions between IT neurons and object recognition behavior, our results guide the development of next-generation biologically constrained brain models by offering novel cell-type specific neural benchmarks.

## Introduction

Distributed neural population activity in the macaque inferior temporal (IT) cortex, which lies at the apex of the visual ventral stream hierarchy, is critical in supporting an array of object recognition behavior^1–3^. Previous research, however, has been largely agnostic to the relevance of specific cell types, inhibitory (Inh) vs. excitatory (Exc), in the formation of “behaviorally sufficient” IT population codes^2^ that can accurately predict primates’ image-by-image object discrimination patterns. In this study, we aim to identify how these distinct classes of neurons in the IT cortex jointly contribute to object recognition behavior.

In the mammalian neocortex, approximately 15-25% of neurons are GABAergic inhibitory neurons, with the majority being glutamatergic excitatory neurons^4–6^. A rich body of research suggests that inhibitory and excitatory neurons play distinct roles in shaping the functional representation of images in the IT cortex. While some studies have revealed a comparable level of stimulus selectivity between these two classes of neurons^7^, others propose a higher degree of stimulus selectivity in excitatory neurons^8^. In addition, local inhibitory circuits are also shown to shape the stimulus selectivity of excitatory neurons in TE^9^. We argue that specific stimuli choices, brain areas, and methods of neuronal recordings can limit the scope of such neuronal characterization without explicitly linking these properties with a specific behavior. Instead, we propose leveraging the behavioral relevance of cell types, across a battery of tasks, to reconcile these diverse findings and motivate an approach to ground the studies of their functional roles (the term “function” relating to relevance in a behavior rather than an interpretable stimulus or neuronal property).

A falsifiable way to generate insights on how diverse neuronal populations in the IT cortex represent visual stimuli and participate in specific behaviors is to develop image-computable artificial neural network (ANN) models of the ventral stream and the behaviors it supports^10^. While ANN models from the family of deep convolutional neural networks explain primate object recognition behavior at unprecedented levels, these models currently present many shortcomings^11–13^. What are the visual system’s most functionally relevant properties lacking in these models? Of note, current ANN models of the ventral stream^14,15^ do not explicitly represent neuronal cell-type. This is in contrast with other attributes of the IT cortex like connectivity patterns^16,17^, functional topography^18–20^, learning mechanisms^21,22^, and hypothesized evolutionary training objectives^23,24^ that has been recently considered to augment models of object recognition. Given the distinct nature of the Exc/Inh ratio in the cortex^4,5^, and the previous reports of differences in their responses to visual stimuli^7,8^, we reasoned that understanding the cell-type specific constraints will be critical to developing more brain-aligned models of object recognition behavior.

In this study, we performed large-scale neural recordings while monkeys fixated images presented in their central field of view. Monkeys also performed 28 different binary object discrimination tasks. We identified putative inhibitory and excitatory neurons in the IT cortex using spike shape-based sorting techniques. We then compared the strength of behavioral predictions of neural decoding (“readout”) models constructed from specific (putative) cell types in the IT cortex. Overall, we observed that decoding strategies derived from excitatory neurons significantly outperform those produced by inhibitory neurons in overall accuracy and image-by-image match to monkey behavioral patterns. Interestingly, we observed that inhibitory and excitatory neurons explained partially overlapping as well as unique fractions of the monkeys’ behavioral variance, emphasizing their distinct roles in object recognition behavior. While evaluating how current ANN explain the response patterns of the specific IT cell types, we observed that all tested ANNs predicted Exc neurons significantly better than Inh neurons. In sum, our results provide correlative evidence for cell-type specificity in the linkage between IT population activity and object recognition behavior, along with the novel cell-type specific constraints for guiding the next generation of brain models.

## Results

As outlined above, we first recorded visually driven neural activity in the macaque IT cortex and categorized them into two separate classes based on their spike waveform shape: narrow spiking (putative inhibitory) and broad spiking (putative excitatory) neurons. Below, we first report the diversity in the response properties of these neuronal cell types and then link these responses to the measured behavioral variances in the macaques. Finally, we evaluate how current ANN models of ventral stream explain these neural responses.

### Identification of putative excitatory and inhibitory neurons

Multiple neurophysiological studies have shown that inhibitory neurons elicit short-duration action potentials^25–28^ which are discharged at higher frequencies. This has led to the term fast spike (FS)^29^ or “thin spikes”^30^ for inhibitory neurons, compared to regular spike^30^ (RS) for excitatory neurons. Though a small portion of inhibitory neurons exhibit broader action potentials^27,31^, and a small fraction of excitatory neurons have narrower action potentials^32,33^, this physiological difference of narrow vs. broad spikes has been widely used to classify extracellularly recorded neurons in several primate neocortical areas, including V4 ^34^, inferior temporal (IT) cortex ^8,35^, and prefrontal cortex (PFC) ^36–39^. Therefore, we leveraged similar spike shape-based criteria (see Methods for details) to classify neurons into putative excitatory and inhibitory cell types. As shown in **Figure 1A**, we recorded neural activity across the macaque IT cortex while the animals passively fixated on images presented for 100ms each. The stimulus set consisted of 640 images spanning 8 objects, with 80 variations per object across changes in position, size, orientation, and background (see Methods). The raw neural data (n=277 units) was then sorted into single neurons using Tridesclous, as made available through the SpikeInterface framework (**Figure 1B**). To ensure high-quality single-unit isolation, we applied three standard quality metrics. First, we computed the inter-spike interval (ISI) violation ratio^40^. Because neurons have a biophysical refractory period (∼1–2ms), spikes occurring within very short intervals (<1.5ms) likely reflect contamination from multiple neurons rather than true physiological firing. Units with higher ISI violation ratios are therefore more likely to represent multi-unit activity. We excluded units with ISI violation ratio > 0.4 (n = 74), leaving 203 units (**Figure 1C**). Second, we evaluated the presence ratio, which quantifies the fraction of the recording session during which a unit is active. Low presence ratios suggest electrode drift or unstable recordings. We excluded units with presence ratio < 0.9 (n = 4), resulting in 199 stable single units (**Figure 1D**). Third, we examined the amplitude cutoff, an estimate of the fraction of spikes falling below the detection threshold (i.e., false negatives). An amplitude cutoff of 0.1 indicates that approximately 10% of spikes are estimated to be missing. After filtering for ISI violations and presence ratio, all remaining units exhibited amplitude cutoff < 0.1 (**Figure S1**), indicating high isolation quality. We then classified neurons based on spike waveform shape using two features: peak-to-valley duration and recovery time (**Figure 1E**). Peak-to-valley duration measures the time between the negative trough and subsequent positive peak of the extracellular waveform (**Figure S1**). Recovery time measures the time from the trough until the waveform returns to baseline (**Figure S1**). Consistent with prior work, putative inhibitory (narrow-spiking) neurons are expected to exhibit shorter peak-to-valley durations and faster recovery times, whereas putative excitatory (broad-spiking) neurons exhibit longer waveform durations. To objectively identify clusters, we first estimated the optimal number of clusters using the Bayesian Information Criterion (BIC), which indicated three clusters (**Figure 1E**). We then fit a Gaussian Mixture Model to classify neurons accordingly. This procedure identified 162 (monkey N: 41, monkey B: 121) broad-spiking neurons (putative excitatory, Exc) and 31 (monkey N: 22, monkey B: 9) narrow-spiking neurons (putative inhibitory, Inh) neurons (**Figure 1F**). Interestingly, as these numbers demonstrate, ∼16% of our pooled neurons across monkeys could be classified as putatively inhibitory, which is consistent with prior reports^5,65,8,35,41,42^.

**Figure 1.**
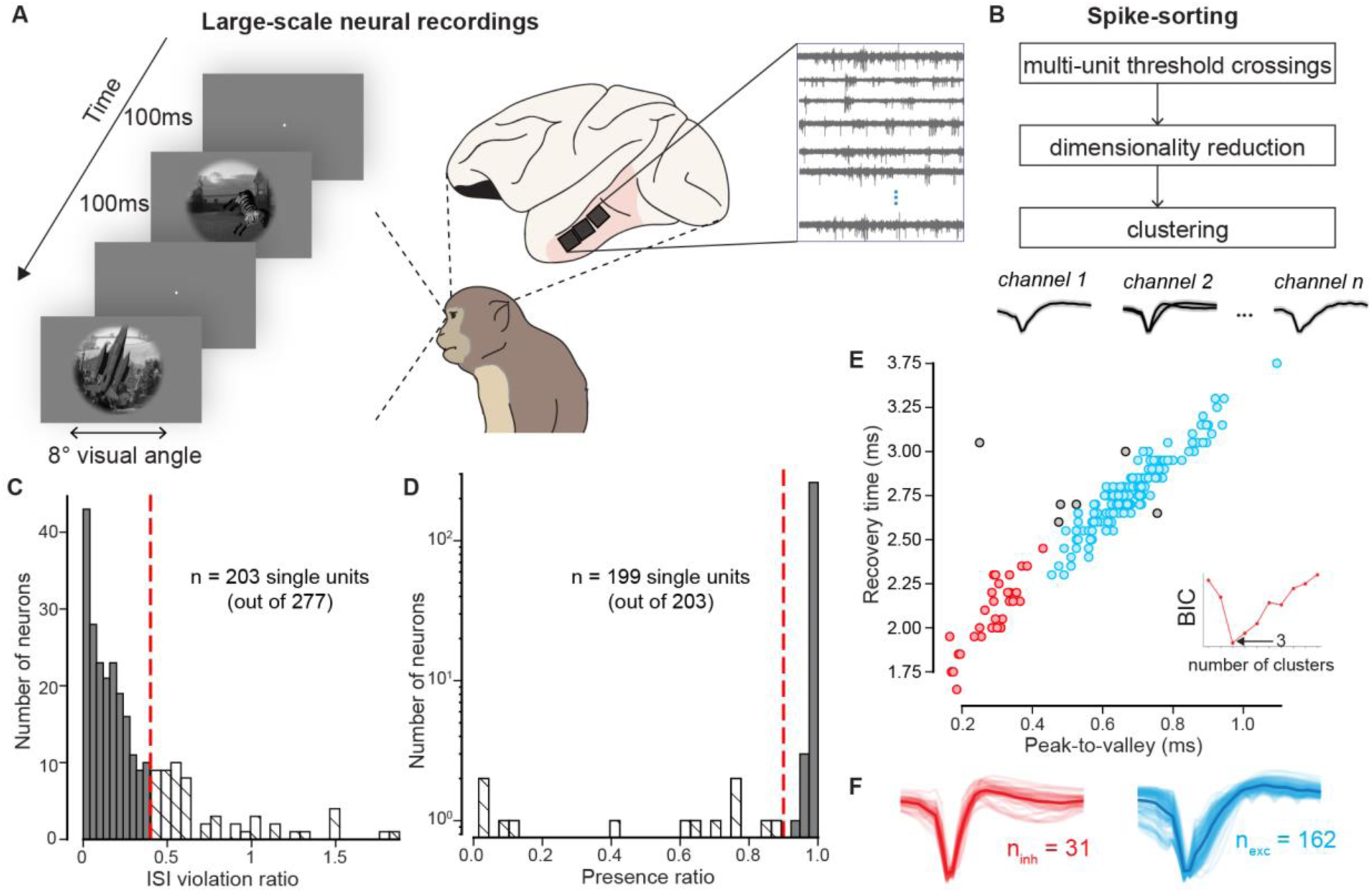
Identification of putative excitatory and inhibitory neurons. **(A)** Neural responses we recorded from inferior temporal (IT) cortex using chronically implanted multi-electrode Utah arrays while the monkeys passively fixated on images (n=640). Inset shows a schematic of bandpass filtered voltages across neural recording sites. (B) Spike-sorting pipeline used to convert neural responses from multi-unit sites into single units and putative cell types. (C) Distribution of ISI violation ratios across neurons. Units with ISI violation ratio higher than 0.4 (n=74) were excluded from further analyses. (D) Distribution of presence ratios across neurons. Units with a presence ratio lower than 0.9 (n=4) were excluded from further analyses. (E) Scatter plot of recovery time versus peak-to-valley waveform duration. Clustering based on these waveform features identified three groups: putative inhibitory neurons (red, n = 31), putative excitatory neurons (blue, n = 162), and excluded units (gray). (F) Illustration of the waveforms for the selected putative inhibitory and excitatory neurons.

### Putative excitatory and inhibitory neurons show distinct functional response properties

To ensure that the experimental noise did not affect the recordings of the Exc and Inh neurons differentially, we first compared the image rank-order response reliabilities (see Methods: Image rank-order response reliability per neuron) of the Inh and Exc neural responses in the 70-170 response window. Although inhibitory neurons exhibited slightly higher reliability than excitatory neurons (ΔReliability = 0.13, unpaired two-sided Wilcoxon test: *z=3.032, p=0.002*) both populations showed relatively high internal consistencies (mean reliabilities = 0.70 and 0.57 for Inh and Exc respectively) indicating that both groups provided robust and reliable stimulus-driven responses (see also **Figure S2** for the distribution of reliability values and the temporal dynamics of reliability). We next examined canonical physiological properties between cell types. We first compared the temporal dynamics of putative excitatory and inhibitory neurons following stimulus onset. **Figure 2A** shows the evolution of firing rate from 0 to 290ms. Inhibitory neurons exhibited consistently higher firing rates throughout the stimulus-driven period. To quantify this difference, we compared baseline (response to grey images within the 70-170ms time window, see Methods: Baseline Firing Rates) and maximum evoked (responses to the stimulus eliciting the strongest response within the same 70–170ms window, see Methods: Maximum Evoked Firing Rates) firing rates of these two neural populations. Consistent with prior reports^8,26^, we observed that inhibitory neurons exhibited significantly higher baseline firing rates (unpaired two-sided Wilcoxon test: Inh>Exc, z=3.961, p<0.001) and higher maximum evoked firing rates (unpaired two-sided Wilcoxon test: Inh>Exc, z=4.534, p<0.001) compared to excitatory neurons (**Figure 2B-C**; Exc: Blue, Inh: Red). We next examined the difference in response latency between the two populations by estimating the response latency based on the evolution of the reliability over time (see Methods: Latency). Specifically, we computed the reliability for each neuron in successive 10ms time bins, from 0 to 290ms after stimulus onset. For each neuron, latency was defined as the first time bin in which reliability reached 0.2 Results were qualitatively similar when using alternative reliability thresholds (see **Figure S2**). **Figure 2D** shows the distribution of latencies for excitatory and inhibitory neurons. Excitatory neurons exhibited significantly longer latencies than inhibitory neurons (unpaired two-sided Wilcoxon test: *z = 5.695, p < 0.001*, mean latencies: Exc = 117ms and Inh = 87ms), consistent with the faster temporal dynamics typically attributed to inhibitory cells^8^. Beyond firing rate differences, we compared response variability across stimuli (see Methods: Response Variance across images). We observed that the response variance is significantly higher in inhibitory neurons (Exc< Inh, unpaired two-sided Wilcoxon test: z=-4.991, p<0.001, normalized histogram shown in **Figure 2E** for both Exc: blue and Inh: red neurons). This finding is similar in nature to broader tuning preferences for stimulus orientation and spatial frequency observed in inhibitory neurons in the mouse^43^ and cat^44^ visual cortex. In contrast, Exc neurons exhibited significantly greater stimulus selectivity (**Figure 2F**, unpaired two-sided t-test: *t(191)=3.193, p=0.002*), quantified using the Depth of Selectivity (DOS) metric^45,46^ (see Methods: Stimulus Selectivity). DOS measures how concentrated a neuron’s responses are across stimuli, with higher values indicating sharper tuning. Importantly, we observed the same qualitative pattern when using alternative selectivity measures, including tuning breadth^47^ (*t(191)=3.341, p=0.002*) and the selectivity index^48^ (*z=2.306, p=0.021*). This result, consistent with prior observations^8^, indicates that excitatory neurons respond more selectively to specific images, whereas inhibitory neurons exhibit broader tuning. Finally, we examined the category selectivity (with each category corresponding to one object, see Methods: Category Selectivity) across multiple temporal windows (**Figure 2G**). Each point represents a time window (marker size reflects bin size; color indicates the end time of the window, see also **Figure 3A**). Across conditions (n=245 windows), excitatory neurons exhibited higher levels of category selectivity compared to inhibitory neurons (paired two-sided Wilcoxon test across conditions: z= 5689.0, p<0.001). This result demonstrates that Exc neurons tend to respond more similarly to images with the same object compared to other objects. Together, these results reveal complementary functional profiles: inhibitory neurons respond earlier and more strongly, with higher overall response variance, whereas excitatory neurons exhibit greater stimulus and category selectivity. These differences raise the possibility that excitatory populations may support more linearly separable object representations.

**Figure 2.**
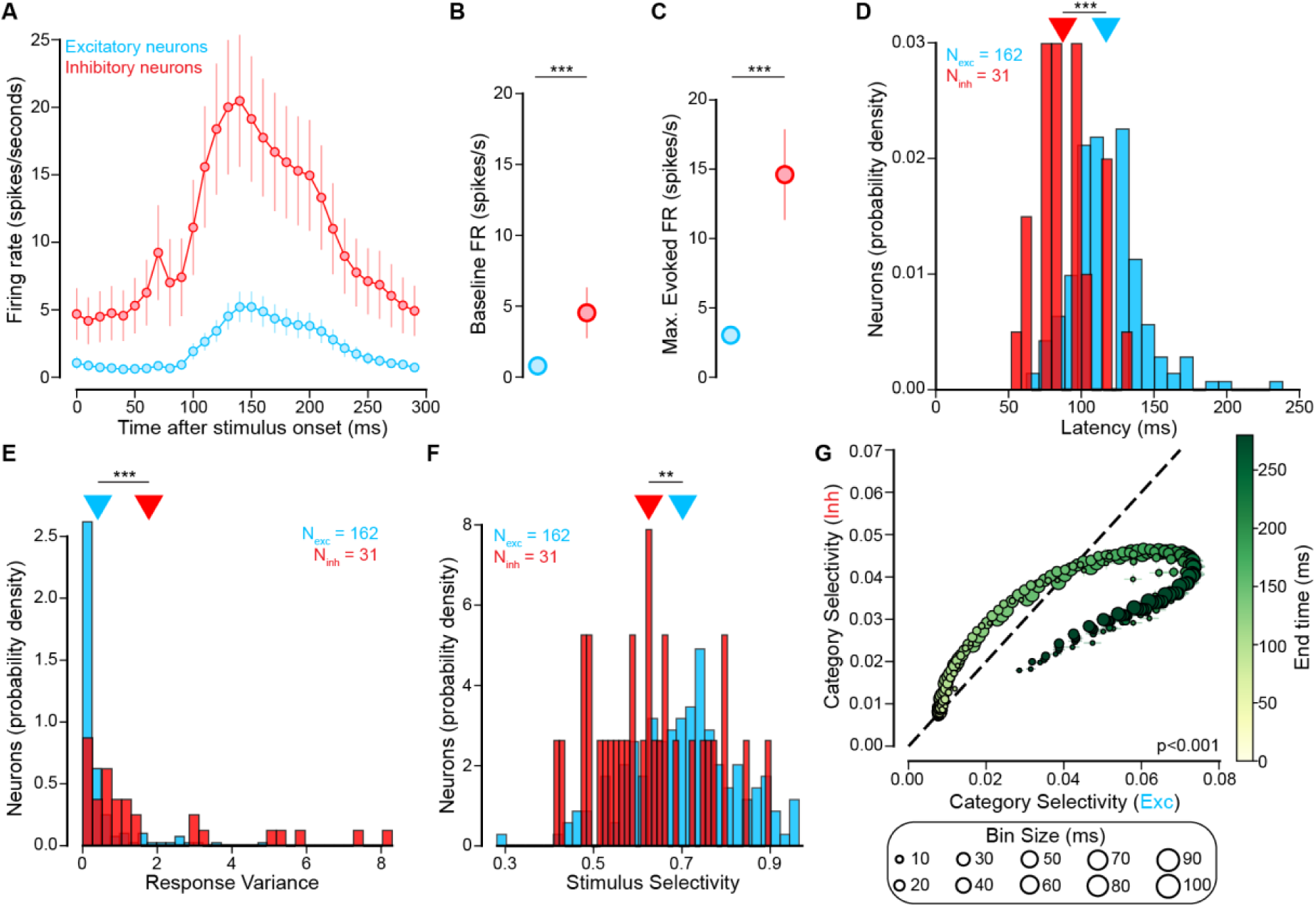
Putative excitatory and inhibitory neurons show distinct functional response properties. (**A**) Evolution of firing rate (mean ± MAD across neurons) over time for Exc (blue) and Inh (red) from 0 to 290ms after stimulus onset. (**B**) Baseline (0-50ms) firing rates (mean ± MAD across neurons) for Exc (blue) and Inh (red) neurons. Unpaired two-tailed Wilcoxon test show higher baseline firing rates for Inh neurons (*z = 3.961, p < 0.001*). (**C**) Max evoked (70-170ms) firing rates (mean ± MAD across neurons) for Exc (blue) and Inh (red) neurons. Unpaired two-tailed Wilcoxon test show higher evoked firing rates for Inh neurons (*z = 4.534, p < 0.001*). (**D**) Histogram of latencies. Latency, computed per neuron, is the time (in ms) at which each neuron’s reliability first reaches 0.2. Unpaired two-tailed Wilcoxon test show higher latency for Exc neurons (*z = 5.695, p < 0.001*). (**E**) Histogram of response variances. Response variance, computed per neuron, is the variance in mean response in 70-170ms interval across all 640 images. Unpaired two-tailed Wilcoxon test show higher variance for Inh neurons (*z = -4.991, p < 0.001*). (**F**) Histogram of stimulus selectivity. Stimulus selectivity, computed per neuron in 70-170ms interval, quantifies how concentrated a neuron’s responses are across all 640 images. Unpaired two-tailed t-test show higher selectivity for Exc neurons (*t(191) = 3.193, p = 0.002*). (**G**) Comparison of category selectivity across conditions between Exc (x-axis) and Inh (y-axis) populations. The diameter of the dots indicates the size of the temporal window; the color indicates the end of the temporal window. Paired two-tailed Wilcoxon test across conditions show higher category selectivity for Exc neurons (*z = 5689.0, p < 0.001*). Statistics were computed across conditions (N = 245). (**A-F**) Statistics were computed across neurons: 162 excitatory neurons and 31 inhibitory neurons. ***: p<0.001, **: p<0.01.

**Figure 3.**
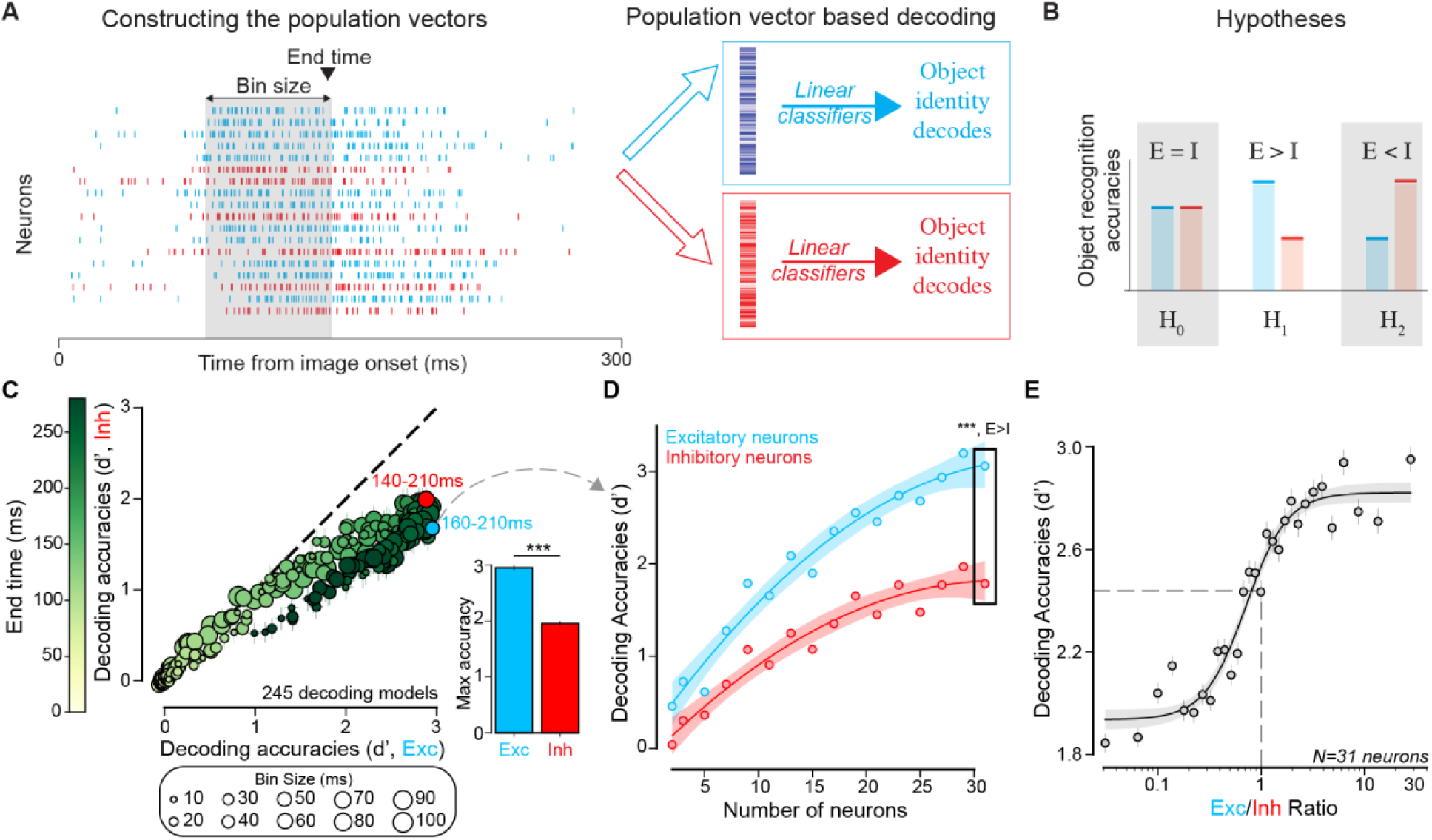
Putative excitatory neurons produce higher accuracy in object discrimination. (**A**) We trained 245 linear decoders (with varying the bin size and end time) on Exc and Inh populations for object recognition. Each decoder was trained on the mean response of neurons in a population. Since the mean response is determined by the number of spikes in some interval characterized by two parameters, namely the end time with respect to stimulus onset and the bin size, we constructed multiple decoders by varying this interval. (**B**) One hypothesis (H_0_) is that linear decodes from both populations would do equally well on object identity discrimination. Another hypothesis (H_1_) is that Exc based decodes would perform better than Inh based decodes; or alternatively (H_2_), Inh based decodes would perform better than Exc based decodes. (**C**) Performance of decoders trained on Exc and Inh populations. Each data point is a linear decoder (n=245) constructed by varying End time and bin size. Inset shows that the best Exc-based decodes (160-210ms) reach higher accuracy than the best Inh-based decodes (140-210ms, two-sided paired Wilcoxon test: *z = 9003.0, p < 0.001*). Error bars represent MAD over random images (n=640). Statistics were computed across images (n=640). (**D**) Performance of the ‘best’ Exc decoder (160-210 ms) as we varied the total number of (Exc vs. Inh) neurons sampled. Error bars represent MAD over reliability-matched sampling of neural population (n=40). Two-sided paired Wilcoxon test across images (n=640) show higher decoding accuracy at n=31 neurons (*z = 2261.0, p < 0.001*) (**E)** Performance of the ‘best’ decoder as we varied the Exc/Inh ratio. Total neurons was always kept at 31. A saturating Hill function provided the best fit (*R² = 0.94*, *RMSE = 0.084*). Error bars represent MAD over reliability-matched sampling of neural population (n=200). (**C-D**) ***: p<0.001

### Putative excitatory neurons produce higher accuracy in object discrimination

As described in the previous sections, the excitatory and inhibitory populations differed in both their number of recorded neurons and intrinsic properties such as response reliability, which could potentially confound population-level decoding comparisons. Therefore, unless otherwise specified, all subsequent analyses were performed on populations matched for both neuron count (using n=31 neurons) and reliability (see Methods: Reliability-matched sampling).

Because Exc neurons exhibited higher object category selectivity (**Figure 2**), we next asked whether this increased selectivity translates into improved object identity decoding. To test this, we constructed population vectors from either excitatory (ticks and population vector in blue; **Figure 3A**) or inhibitory neurons (ticks and population vector in red; **Figure 3A**) and trained linear classifiers to discriminate object identities (see Methods: Estimation of IT population decode accuracies). We formalized three competing hypotheses. The first possibility is that Exc could lead to higher accuracies on an object discrimination task compared to Inh neurons (H_1_, **Figure 3B**). The other possibilities are that Inh neuron-based population decodes outperform the Exc based ones (H_2_, **Figure 3B**), and that these two populations produce equally accurate decoders (H_0_, **Figure 3B**). In these analyses, spike counts were computed within a temporal integration window defined by two parameters: window end time and bin size (**Figure 3A**). By systematically varying these parameters, we constructed 245 linear decoding models per population, spanning a broad range of temporal windows and integration widths. Across this parameter space, we observed that the decoders trained on excitatory populations consistently achieved higher discrimination performance (measured in d’) than those trained on inhibitory populations (rejecting H_0_ and H_2_, **Figure 3C**, two-sided paired Wilcoxon test across conditions: *z = 1701, p < 0.001*). Additionally, the strongest decoders Exc-based decoders (integration window 160 to 210 ms from image onset, mean d’=2.95±0.84 s.e.m) significantly outperformed the strongest Inh-based decoders (140-210ms, mean d’=1.96±0.78 s.e.m, inset **Figure 3C**, two-sided paired Wilcoxon test across images: *z = 9003.0, p < 0.001,* see also **Figure S3** for results per monkey). Thus, although both populations carry linearly decodable object information, excitatory representations yield systematically higher decoding accuracy. Because Exc neurons exhibited higher object selectivity (**Figure 2F**), we performed an additional selectivity-matched sampling analysis in which excitatory neurons were subsampled to match the inhibitory population’s DOS distribution (**Figure S4A**). Even under this constraint, excitatory populations continued to significantly outperform inhibitory populations in decoding performance (**Figure S4B**), indicating that the advantage cannot be explained solely by differences in stimulus selectivity. However, the accuracies when measured in this fashion depend heavily on the number of neurons sampled. Therefore, we sampled from 2 to 31 neurons (randomly for both Exc and Inh pools, given the 160-210ms integration window). For both populations, accuracy increased with the number of neurons (**Figure 3D**). Additionally, Exc-based decoders consistently outperformed Inh-based decoders across sampled population sizes (at n=31 neurons, Exc>Inh, two-sided paired Wilcoxon test across images: *z = 2261.0, p < 0.001*), and the difference in accuracies between the two groups increased as a function of the number of neurons (at n=5, ΔExc-Inh=0.25, vs. at n=31, ΔExc-Inh=1.28). This scaling analysis suggests that the observed difference is not driven by small-sample variability and likely represents a lower bound on the true population-level difference. Importantly, while Exc outperformed Inh based decodes, it is important to note that the linearly separable category information contained in the representation of the Inh neurons is significantly greater than zero (with a best d’ ∼2) and only 33.68% (mean) lower than the Exc neurons. Therefore, we further probed how the decoding accuracies evolve as we vary the ratio of participation of the Exc and Inh neurons in the neural population pool (keeping the overall number of neurons constant at n=30). Our results (**Figure 3E**) show that decoding accuracies monotonically increase as a function of the Exc/Inh ratio with the inclusion of more Exc neurons. To further characterize how each population contributes to decoding performance, we performed an additional nested analysis in which one population was held fixed while neurons from the other population were incrementally added at matched total neuron count. Across both complementary analyses, adding excitatory neurons consistently produced larger improvements in decoding accuracy than adding inhibitory neurons, and at no population size did the inclusion of inhibitory neurons exceed the performance achieved by additional excitatory neurons (see **Figure S4C-D**). Together, these findings indicate that excitatory populations provide a more behaviorally relevant representation of object identity, motivating us to next test how each population’s responses align with image-by-image behavioral performance.

### Putative excitatory neuron based decodes produce higher image-level consistency with monkeys’ object discrimination behavior

Although the decoding accuracies estimated above quantify the amount of linearly separable object information present in the Exc and Inh neural populations, high decoding accuracy alone does not guarantee behavioral relevance. We therefore asked whether population decoders derived from each cell type predict monkeys’ image-by-image object discrimination behavior. We first measured the behavioral responses of the same 2 macaques on a delayed match-to-sample object discrimination tasks. On each trial, monkeys viewed a Test image for 100ms (containing a target object; one among 8), followed by a choice screen containing canonical views of two objects (the target and a distractor object, see **Figure S5A-B**). The monkeys had to indicate their choice of the target object with a saccade to one of the options (**Figure 4A**). In total, each of the 8 objects served as the target against each of the remaining 7 objects as distractors, resulting in 56 distinct target–distractor discrimination conditions. We then estimated an image-level metric (referred to as **B.I_1_**) from the measured behavioral performances. In brief, the **B.I_1_** metric reflects the performance of the monkey in identifying the target object in each image, when tested across all possible distractor categories (see Methods: Behavioral Metrics and **Figure S5C**, also for more detail see the estimation of **B.I_1_** in^11^). The trial level split-half reliability of this metric, after behavioral trials were pooled across 2 monkeys (yielding ∼240 repetitions per image) was 0.93 ± 0.004 (mean ± SD). We then quantify the consistency of the neural population-based predictions (**N.I_1_**) with the behaviorally estimated **B.I_1_** as the trial-level noise corrected correlation between the two (see Methods: Consistency of **N.I_1_** with **B.I_1_)**. Each point in **Figure 4C** corresponds to one linear decoder (n=115, we excluded the integration window leading to low reliability of the **N.I_1_**) defined by a specific temporal integration window (bin size and end time; see **Figure 3A**). Across temporal windows, Exc-based decoders consistently achieved higher behavioral consistency that Inh-based decoders (**Figure 4C**, two-sided paired Wilcoxon test across conditions: *z = 420.0, p < 0.001*, rejecting H_0_ and H_2_, **Figure 4B**). The highest consistency Exc-based decoders (best consistency reached at 70-150ms after stimulus onset, mean=0.63 ± 0.01 MAD) significantly outperformed the highest-consistency Inh decoder (110-120ms, mean 0.52 ± 0.02 MAD, inset **Figure 4C**, permutation test across resampling of the neural population: *D = 0.112, p < 0.001,* see also **Figure S6A-C** for results per monkey). Importantly, this difference in behavioral consistency was not driven by differences in overall decoding accuracy. When we performed an additional control analysis in which excitatory and inhibitory populations were subsampled to match decoding accuracy prior to computing behavioral consistency, Exc-based decoders continued to exhibit significantly higher consistency with behavior (**Figure S6D-E**). These results indicate that the greater behavioral alignment of excitatory populations cannot be explained solely by their higher object classification performance.

**Figure 4.**
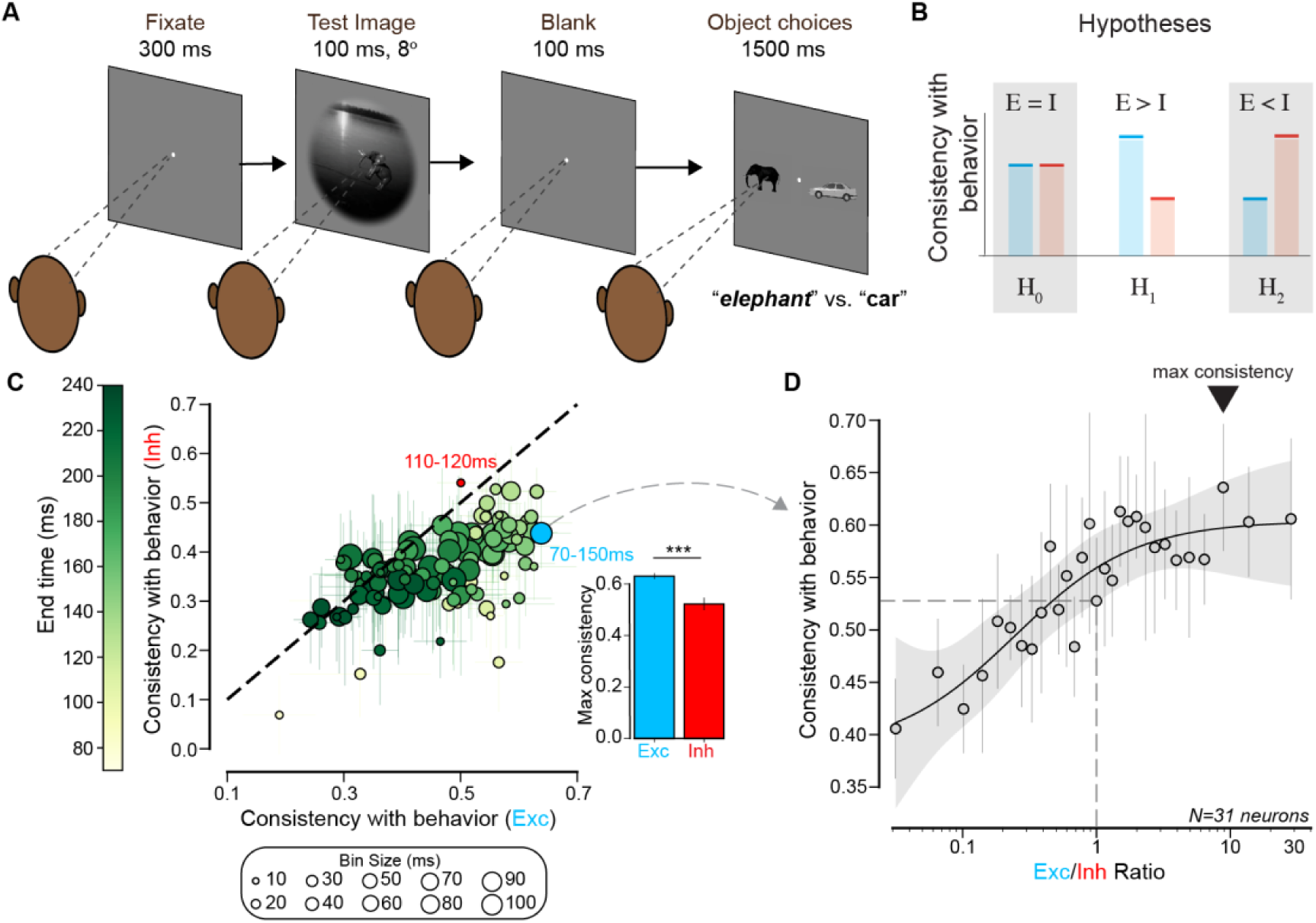
Putative excitatory neuron based decodes produce higher image-level consistency with monkeys’ object discrimination behavior. **(A)** We tested 2 monkeys on multiple delayed match to sample binary object discrimination tasks. Monkeys initiated a trial by fixating on a central white dot for 300 ms, following which a test image (out of 640 images) containing the target object (shown elephant) was presented in the central 8°. This was followed by a blank screen for 100 ms, and then canonical views of the target (elephant) and distractor (car; one out of 7) object were shown on the screen (randomly assigned distractor for each trial). The monkeys had 1500ms to freely view the object choices and indicate their selection of the target by fixating on it for 400 ms. (**B**) One hypothesis (H_0_) is that linear decodes from both populations would be equally consistent with monkey behavior. Another hypothesis (H_1_) is that Exc based decodes would be more consistent with monkey behavior; or alternatively (H_2_), Inh based decodes would be more consistent with monkey behavior. The green rectangle indicates the hypotheses that the data couldn’t falsify. (**C**) Behavioral consistency of Exc and Inh based decodes. Each data point is a linear decoder (n=115) based on a specific choice of bin size and end time (refer Figure 3A). Inset shows that the Exc-based decodes highest consistency (70-150ms) outperformed the highest consistency from Inh-based decodes (110-120ms, permutation test: *D = 0.112, p < 0.001*). Statistics were computed using permutation tests across reliability-matched samplings of 30 neurons from the Exc and Inh sets respectively (n=20). Error bars represent MAD across random draws (n=20). The highest consistency readout was from Exc neural response in 70 to 150ms interval. ***: p<0.001. (**D**) Behavioral consistency of the highest consistency decoder found in (**C**) for varying Exc/Inh ratio (with a fixed number of neurons. Max behavioral consistency is achieved for E/I ratio ∼6-7. A saturating Hill function provided the best fit (*R² = 0.81*, *RMSE = 0.027*). Error bars denote reliability-matched sampling of neurons to construct the neural pool (n=1000).

Given that the inhibitory neurons represent a significant fraction of object information and that these decodes correlate highly with our measured behavioral metrics, we further investigated whether combining Exc and Inh neurons in the population vector produce more behaviorally aligned decodes. Specifically, we varied the Exc/Inh ratio while keeping the total population size fixed (**Figure 4D**). In sharp contrast to the decoding accuracies (**Figure 3E**), where adding Exc neurons always increased decoding accuracies (tail slope for E/I ratio ≥3: mean=0.076 95% CI [0.045, 0.108]), behavioral consistency did not exhibit continued improvement at higher ratios (**Figure 4D**). While consistency increased from low to moderate Exc/Inh ratio, slope estimates at higher ratios (E/I ratio ≥3) were not statistically different from zero (mean=0.0038 CI [−0.185, 0.236]). Across the entire range from E/I ratio = 3 to E/I ratio = 31, the mean change in consistency was less than 6% (Δconsistency=0.0267), indicating minimal additional behavioral gain despite substantial increases in excitatory proportion. Thus, whereas increasing the proportion of excitatory neurons continuously improves linearly decodable object information, behavioral alignment shows diminishing returns once excitatory neurons dominate the population. This plateau indicates that maximally excitatory populations already achieve similarly high behavioral consistency. We next examine whether excitatory and inhibitory populations nevertheless explain overlapping or distinct components of behavioral variance.

### Putative excitatory and inhibitory neurons explain non-overlapping variances in object discrimination behavior

To quantify how much variance in object discrimination behavior is uniquely explained by Inh and Exc populations, we performed a split-half partial correlation analysis (see Methods: Partial correlation of **N.I_1_** with **B.I_1_**). Specifically, for each population, neural responses were divided into two independent halves along the repetitions dimension. For a given temporal window (defined by end time and bin size, see **Figure 3A**), we computed the image-level neural prediction (**N.I_1_**) separately for each half. We then correlated the first half of one population with behavioral I_1_ while controlling either (i) for the second half of the same population (to account for internal reliability and shared noise) or (ii) for the first half of the other population (to estimate unique cross-population contributions). This procedure, repeated multiple times, ensured independence between the neural predictor and the variable being controlled for. **Figure 5A** shows the behavioral consistency of Exc-based decoders across temporal windows (n=245) when controlling for the split-halves (x-axis) against the correlation when controlling for inhibitory readouts (y-axis, see **Figure S7** for partial correlation vs. accuracy). Points falling below the unity line indicate that part of the behavioral variance explained by Exc neurons overlaps with inhibitory representations. However, most points remain well above zero, indicating that a substantial fraction of behavioral variance is uniquely captured by Exc neurons even after accounting for Inh activity (paired two-tailed t-test across conditions: *t(244) = -7.69, p < 0.001*). The temporal window yielding the strongest behavioral alignment (from **Figure 4C**, 70–150 ms; star) retains robust partial correlation under cross-population control. **Figure 5B** shows the complementary analysis for inhibitory neurons. Here, the x-axis reflects behavioral correlation controlling for inhibitory split-half reliability, and the y-axis reflects correlation controlling for excitatory readouts. While inhibitory neurons retain non-zero unique behavioral variance (paired two-tailed Wilcoxon test across conditions: *z = 916.0, p < 0.001*), many points lie closer to the diagonal and overall magnitudes are smaller, indicating that a larger fraction of inhibitory behavioral variance overlaps with excitatory representations (see also **Figure S7** for similar results per monkey). Direct comparison of the cell-type–specific unique behavioral contributions (**Figure 5C**) confirms that excitatory neurons account for a greater amount of variance uniquely aligned with behavior (paired two-tailed Wilcoxon test across conditions: *z = 1458.0, p < 0.001*). We observed that the Exc cell-type exclusive behavioral variances peaked between 180-280ms. Nonetheless, inhibitory neurons contribute measurable non-overlapping variance across time.

**Figure 5.**
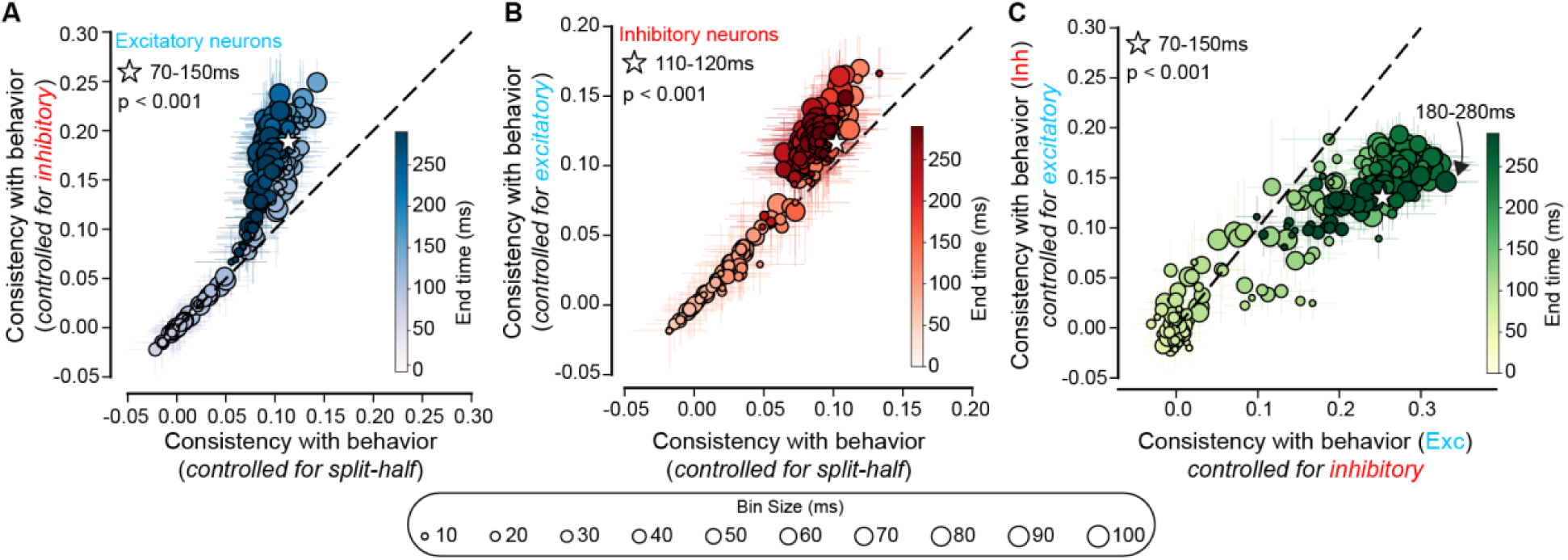
Putative excitatory and inhibitory neurons explain non-overlapping variances in object discrimination behavior. (**A**) Correlating monkey behavior and Exc based decodes (varying bin size and end time, n=245) controlling for readouts from Inh neurons vs. controlling for split-half of Exc decodes. The star indicates the temporal window leading to the highest consistency with behavior (**Figure 4C**). A two-sided paired t-test shows that controlling for Inh **N.I_1_** leads to higher consistency compared to controlling for Exc split-half **N.I_1_** (*t(244)=-7.69, p<0.001*). (**B**) Correlating monkey behavior I_1_, and Inh neurons based decodes (varying bin size and end time, n=245) controlling for readouts from Exc neurons vs. controlling for split-half of Inh decodes. The star indicates the temporal window leading to the highest consistency with behavior (**Figure 4C**). A two-sided paired Wilcoxon test shows that controlling for Exc **N.I_1_** leads to higher consistency compared to controlling for Inh split-half **N.I_1_** (*z=916.0, p<0.001*). (**C**) Comparing correlation of monkey behavior with cell-type unique variances (Exc vs. Inh). Excitatory neurons, after controlling for the Inhibitory population explains more of the measured behavior compared to Inhibitory neurons controlled for the Excitatory population (two-sided paired Wilcoxon test: *z=1458.0, p<0.001*). The star indicates the temporal window leading to the highest consistency with behavior (**Figure 4C**). (**A-C**) Error bars represent MAD over reliability-matched sampling of neural population (n=100). Statistics were computed across conditions (n=245).

Together, these results indicate that excitatory and inhibitory populations provide partially complementary representations of object identity. Excitatory neurons dominate behaviorally aligned variance, but inhibitory neurons contribute additional components not fully redundant with excitatory activity, suggesting that downstream readout mechanisms likely integrate information from both cell types. To understand the origin of these complementary contributions, we next characterize the representational geometry of excitatory and inhibitory populations, examining how their object manifolds differ in structure and separability.

### Putative excitatory and inhibitory populations exhibit distinct representational geometries

To determine whether Exc and Inh populations encode similar object representations, we first quantified manifold geometry^49,50^ of object representations in each population within the 70-170 time window (**Figure 6B**, see also **Figure S8** for best accuracy and best consistency time windows). Effective dimensionality^51,52^, measured by participation ratio (see Methods: Object manifold geometry metrics), was higher in Exc neurons (mean = 6.35 ± 0.67 MAD) compared to Inh neurons (mean = 5.75 ± 0.34 MAD; permutation test across 50 samplings of neurons: *D = 0.896, p < 0.001*). Manifold radius was smaller for Exc populations (mean = 0.34 ± 0.024 MAD) relative to Inh populations (mean = 0.58 ± 0.03; *D = -0.249, p < 0.001*). Manifold capacity was correspondingly greater for Exc neurons (mean = 1.66 ± 0.21 MAD) than for Inh neurons (mean = 0.72 ± 0.07 MAD; *D = 0.959, p < 0.001*). Additionally, distances between category manifolds were larger in the Exc population (mean = 0.61 ± 0.08 MAD) compared to Inh (mean = 0.55 ± 0.05 MAD; *D = 0.101, p < 0.001*). These geometric metrics indicate that category representations in the Exc population occupy a lower-radius, higher-capacity region of neural space with greater inter-manifold separation relative to Inh populations as illustrated in **Figure 6A**. To identify statistical factors contributing to these geometric differences, we quantified response variability across and within objects (70-170ms, **Figures 6C–D**, see **Figure S9** for best accuracy and best consistency time windows, and Methods: Across and within object variance). Across-object variance (the variance of mean firing rate across the 8 objects) was higher for Inh neurons (median = 0.065) than for Exc neurons (median = 0.01; unpaired two-tailed Wilcoxon test: *z = −3.849, p < 0.001*, **Figure 6C**). Within-object variance (variance across 80 images per category, averaged across categories) was also higher in Inh neurons (median = 0.77) compared to Exc neurons (median = 0.10; unpaired two-tailed Wilcoxon test: *z = −5.096, p < 0.001*, **Figure 6D**). Across-object variance contributes to separation between object centroids, whereas within-object variance determines manifold radius. The substantially higher within-object variance in inhibitory neurons expands manifold radius and increases overlap between objects, consistent with the larger radius and lower manifold capacity observed for Inh populations in **Figure 6B**. In contrast, the lower within-object variance in Exc neurons supports more compact and separable object manifolds. Finally, we quantified shared variability via mean pairwise neuron–neuron correlations (see Methods: Inter-neuron correlations) within each population within the 70-170ms time window (**Figure 6E**, see **Figure S9** for best accuracy and best consistency time windows). Inhibitory neurons exhibited higher inter-neuron correlations (mean = 0.073 ± 0.007 MA) than excitatory neurons (mean = 0.045 ± 0.002; unpaired two-tailed t-test: *t(191) = -3.41, p < 0.001*). Increased inter-neuron correlations constrain the number of independent activity dimensions within the population, consistent with the lower effective dimensionality observed for inhibitory neurons.

**Figure 6.**
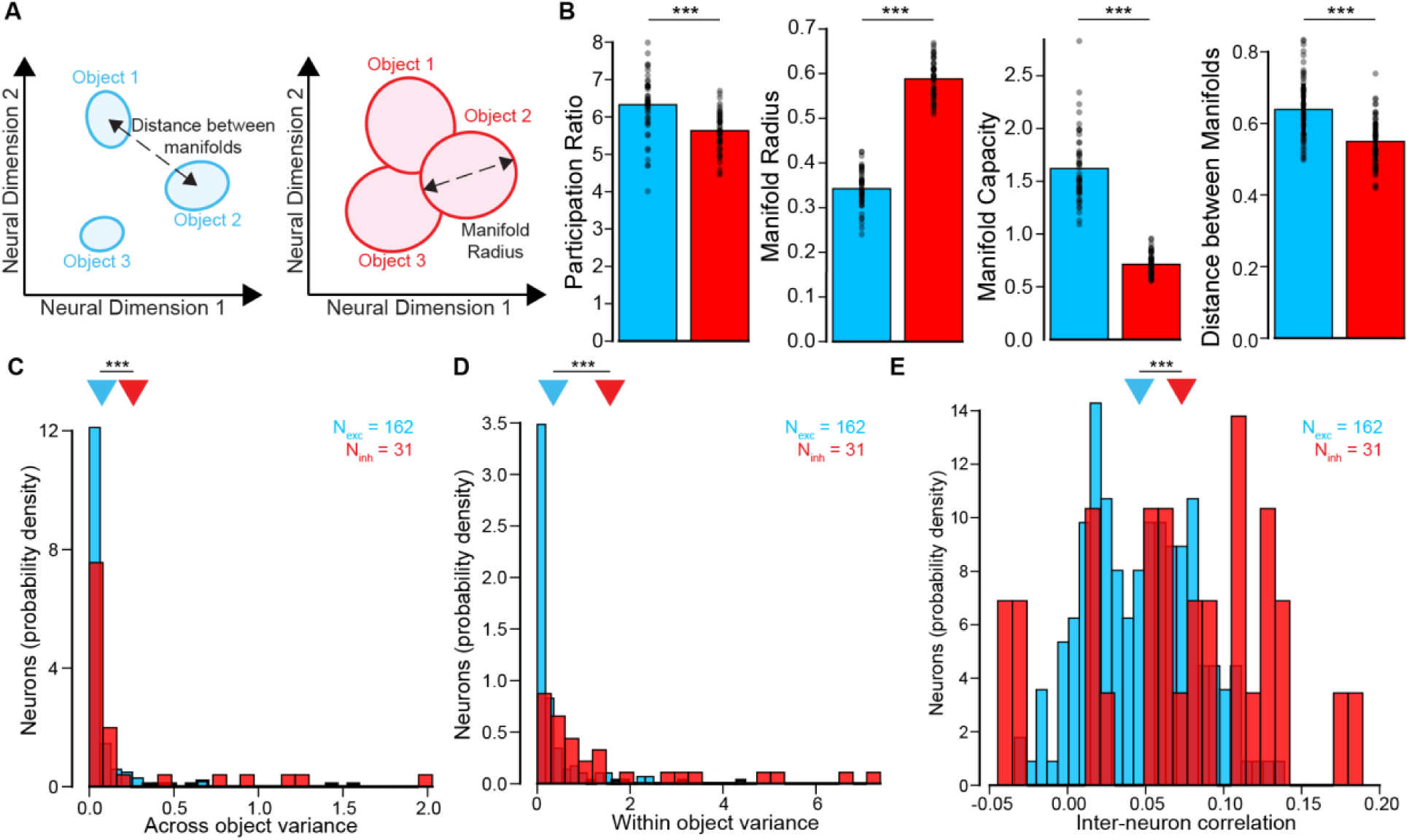
Putative excitatory and inhibitory populations exhibit distinct representational geometries. (**A**) Schematic illustration of representational geometry differences. Excitatory neurons form compact, well-separated category clusters, whereas inhibitory neurons exhibit broader and more overlapping category manifolds. (**B**) Summary metrics of manifold geometry, including participation ratio (effective dimensionality), manifold radius, manifold capacity and distances between manifolds. Each dot represents one sampling of the neural data. (n=50). Statistics were computed using permutation tests across reliability-matched samplings of neural data (***p < 0.001). (**C**) Distribution of across-category variance per neuron (excitatory: blue; inhibitory: red). Across-category variance was computed as the variance of the mean firing rate (70–170 ms post-stimulus) across the 8 object categories. Inhibitory neurons exhibit higher across-category variance than excitatory neurons (unpaired two-tailed Wilcoxon test: *z = −3.85, p < 0.001*). (**D**) Distribution of within-category variance per neuron. Within-category variance was computed as the variance of mean responses (70–170 ms) across the 80 images belonging to each category, then averaged across categories. Inhibitory neurons show higher within-category variance than excitatory neurons (unpaired two-tailed Wilcoxon test: *z = −5.10, p < 0.001*). (**E**) Distribution of inter-neuron correlation per neuron. Inter-neuron correlation was computed as averaged image-level correlation between one neuron and all the others within the population (70–170 ms post-stimulus). Inhibitory neurons exhibit higher inter-neuron correlation than excitatory neurons (unpaired two-tailed t-test: *t(191) = -3.41, p < 0.001*).

Taken together, these analyses demonstrate that Exc and Inh populations differ in both representational similarity and geometric organization. Exc populations exhibit higher manifold capacity, smaller manifold radius, and greater inter-manifold separation, whereas Inh populations show increased within-category variance and stronger shared correlations. These geometric and statistical differences provide a quantitative account of the differential decoding performance and behavioral alignment observed in previous sections. We next test whether contemporary deep neural network models differentially capture these cell-type–specific representational properties.

### ANN models of object recognition better predict putative excitatory than inhibitory neurons

Our results suggest that putative excitatory and inhibitory neurons differ in behavioral alignment (**Figure 3**) and representational geometry (**Figure 5**). Therefore, we next asked whether current ANN models of the ventral stream predict Exc and Inh neurons equally well. To test this, we first tested a state-of-the-art brain-aligned model of the ventral stream, CORnet-S^15^ that has a fixed, committed mapping between macaque ventral stream areas (V1, V2, V4, and IT) and model layers.

We then used linear regression to map the features of the ANN-IT layers onto the responses of individual macaque-IT neurons^16,22^ (see Methods: Prediction of neural responses from ANN features). We quantified predictivity as percent explained variance (%EV) across neurons. Interestingly, we observed that the CORnet-S IT layer tends to predict Exc neurons better than Inh neurons (**Figure 6A**, median %EV_Exc = 52.76 ± 1.06 MAD; mean %EV_Inh = 46.65 ± 2.54 MAD; non-significant two-tailed unpaired Wilcoxon test, *z=0.670, p=0.251*). We also observed similar trends across all tested layers of CORnet-S (dashed bars, V1: *z=-3.97, p=0.654,* V2: *z=-0.309, p=0.621,* V4: *z=0.018, p=0.493*). To further test the generalizability of this result across other ANN models, we next evaluated a panel of 38 high-performing ANN models (e.g., AlexNet^53^, CORnet-S^15^, ResNets^54^ etc.; see Methods: Model selection, **Figure S10**) from Brain-Score^14^ as well as 7 ConvRNNs^85^. Using Brain-Score’s^14^ public dataset, we first established a fixed mapping between a specific model layer (see Methods: Layer selection and **Figure S10**) and the macaque IT. We then quantified representational similarity using noise-corrected CKA between model IT layers and neural responses separately for Exc and Inh populations (**Figure 7B**, see Methods: CKA between ANN features and neural responses). Across models, CKA values were consistently higher for Exc neurons than for Inh neurons (mean CKA_Exc = 0.36 ± 0.036 MAD; mean CKA_Inh = 0.29 ± 0.02 MAD; two-tailed paired Wilcoxon test across models: *z =3.0, p < 0.001*), indicating that model representational structure aligns more closely with excitatory population. We also quantified forward predictivity^22^, mapping model features onto neural responses (**Figure 7C**). Across models, Exc neurons exhibited significantly higher explained variance than Inh neurons (mean %EV_Exc = 48.79 ± 3.29 MAD; mean %EV_Inh = 42.86 ± 3.33 MAD; two-tailed paired Wilcoxon test across models: *z = 0.0, p < 0.001*). Finally, we evaluated reverse predictivity^92^, mapping neural responses onto model features (**Figure 7D**, see Methods: Prediction of ANN features from neural responses). Exc neurons predicted model units significantly better than Inh neurons (mean %EV_Exc = 2.81 ± 1.06; mean %EV_Inh = 2.09 ± 0.76; two-tailed paired Wilcoxon test across models: *z =0.0, p < 0.001*). The weaker reverse predictivity for inhibitory neurons suggests that model feature spaces do not fully capture the structure of inhibitory population activity, even when neural responses are used to predict model units directly. Together, these results demonstrate that ANN models better align with excitatory neurons across multiple representational metrics (CKA, forward predictivity, reverse predictivity) and across diverse architectures (see also **Figure S11** for results per model, **Figure S12** for results per individual monkey and **Figure S13A-C** for results on the 180-280ms time window corresponding to the Exc cell-type exclusive behavioral variance peak). This systematic gap suggests that current ANN models capture representational structure characteristic of excitatory populations more faithfully than that of inhibitory populations. Importantly, this Exc–Inh predictivity difference cannot be trivially attributed to differences in explainable variance: all predictivity values are noise-corrected by each neuron’s response reliability (i.e., normalized by its noise ceiling), and inhibitory neurons in fact exhibit equal or higher reliability and explainable variance than excitatory neurons (see **Figure S2D**), ruling out ceiling effects as an explanation for the observed gap. We next ask whether this differential neural predictivity translates into differences in behavioral predictivity across cell types.

**Figure 7.**
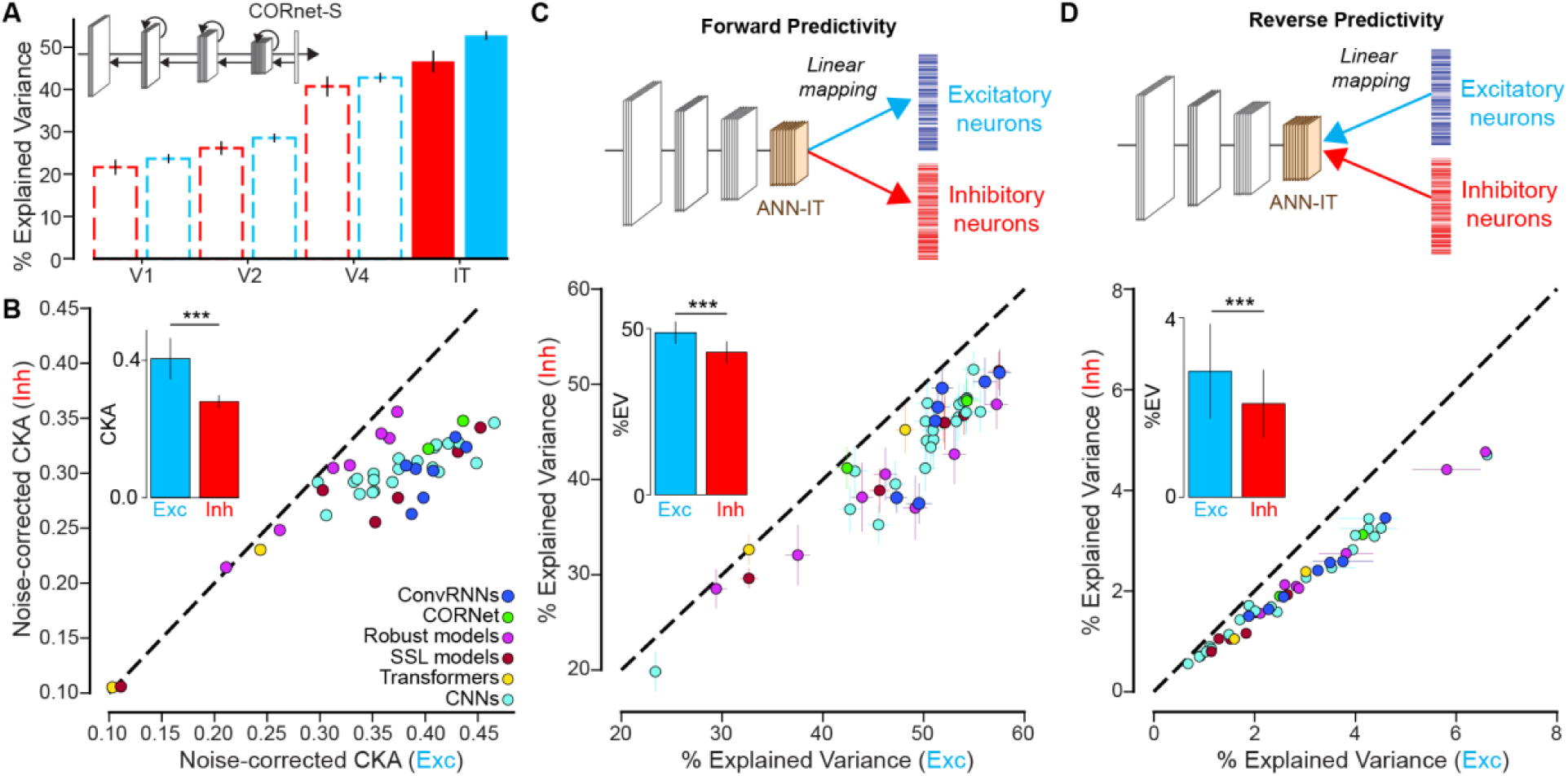
ANN models of object recognition better predict putative excitatory than inhibitory neurons. **(A)** The IT layer of CORnet-S, a model designed to match the ventral visual stream hierarchy (V1, V2, V4, and IT), tends to predict Exc IT neurons better than the Inh neurons. Error bars denote MAD across neurons. **(B)** CKA between models ITs and neural responses (Exc vs. Inh) Each point represents a model. Inset shows mean CKA for excitatory and inhibitory populations. **(C)** Forward predictivity for excitatory and inhibitory neurons across models. Inset shows mean explained variance across models with Exc neurons better predicted that Inh neuron. Error bars denote MAD across neurons. **(D)** Reverse predictivity for excitatory and inhibitory neurons across models. Inset shows mean explained variance across models with Exc neurons predicting model units better than Inh neurons. Error bars denote MAD across model units. **(B-D)** Two-sided paired Wilcoxon test across n=45 models; ***: p < 0.001.

Figure 8A demzonstrates the intuition behind a variance partitioning analysis to address this question. The gray shaded region in the left panel of Figure 8A refers to the behavioral variance explained by the ANNs. In our study such estimates are performed by a correlation between ANN-based I_1_ predictions and the measured behavioral **B.I_1_** vector. Next the green region in the mid panel refers to the behavioral variance explained by the ANNs controlling for a neural decode based variance (e.g., from Exc or Inh neural population decodes). Such an estimate is retrieved by performing a partial correlation analysis between ANN I_1_ and Behavior **B.I_1_** with neural **N.I**_1_ as a control variable. Finally, the right panel is the result of subtracting the first two estimate and provides the image-level behavioral variance explained by ANNs that is common with the neurally explained variance. Figure 8B shows the result of this analysis across 45 models (see **Figure S13** for the plots corresponding to each step). We observe that indeed, consistently across all models, the Exc predicted variance is better captured by ANNs compared to Inh predicted variance (two-sided paired Wilcoxon test across models: *z=0.0, p<0.001*).

**Figure 8.**
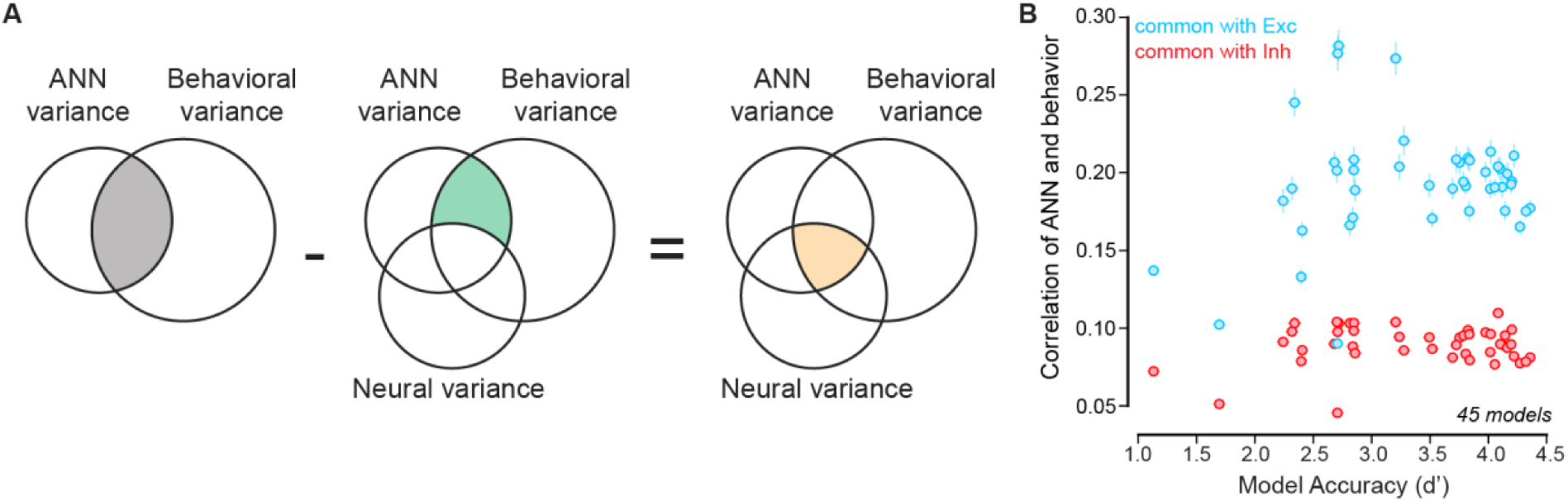
Variance partitioning analysis to estimate the amount of behavioral variance jointly explained by neurons (Exc vs. Inh) and ANN-IT layers. **(A)** Venn diagrams showing shared variance. Left panel: The gray shaded region refers to the behavioral variance explained by the ANNs. Mid panel: The green region refers to the behavioral variance explained by the ANNs controlling for a neural decode based variance. Right panel: The yellow region refers to the result of subtracting the first two estimates and provides the image-level behavioral variance explained by ANNs that is common with the neurally explained variance. (**B)** Comparison of correlation of ANN predictions with behavior that is common with predictions produced by Exc (blue) and Inh (red) neurons (across 45 models).

## Discussions

The primary goal of this study was to probe the cell type specificity of the population activity of the macaque inferior temporal cortex that supports core object recognition behavior. Our results provide evidence that both putative inhibitory and excitatory neurons each make unique contributions to shaping the monkeys’ behavioral reports. Below we discuss our results in the context of their broader implications.

### Limitations related to inhibitory population size

A limitation of the present study is the relatively smaller number of putative inhibitory neurons compared to excitatory neurons. This imbalance reflects both biological reality — inhibitory neurons constitute a minority of cortical populations^5,8,35,41,42^ — and the constraints of extracellular recording and waveform-based classification. Because decoding performance scales with population size, comparisons between cell types could in principle be biased by unequal sampling. To mitigate this concern, all population-level analyses were performed using reliability-matched subsampling procedures, with decoding and behavioral consistency computed from fixed-size populations (n = 30 neurons) repeatedly resampled across iterations. Thus, the reported differences do not reflect trivial scaling with neuron count. Nevertheless, the inhibitory population was drawn from a limited pool, and the resampling procedure (sampling 30 out of 31 inhibitory neurons) necessarily results in substantial overlap across iterations. While this approach provides stable point estimates of decoding accuracy and behavioral alignment, it limits our ability to fully characterize the diversity of inhibitory representations in the broader IT population. In particular, rare inhibitory subtypes or functionally specialized interneurons may be underrepresented. Therefore, our conclusions should be interpreted as describing the recorded inhibitory population rather than the complete inhibitory circuitry of inferior temporal cortex.

Importantly, several aspects of the results argue against the observed differences being an artifact of limited inhibitory sampling. First, inhibitory-based decoders reliably carried significant object information above chance. Second, inhibitory populations explained a unique fraction of behavioral variance even after controlling for excitatory predictions. Third, geometric differences in manifold radius and separability were consistent across temporal windows. Together, these converging findings suggest that the qualitative differences between excitatory and inhibitory populations are robust, even if the precise quantitative magnitude may be refined by larger future datasets.

Future work using higher-density recordings, cell-type–specific tagging, or large-scale calcium imaging could provide more comprehensive sampling of inhibitory subtypes and allow finer-grained comparisons across interneuron classes. Such approaches will be essential to determine whether the representational properties reported here generalize across inhibitory subclasses or are driven by specific functional types.

### Exc/Inh contributions to linking propositions and downstream readout mechanisms

We demonstrate that Exc and Inh neurons provide unique contributions to the object identity solutions. In particular, linear decoders trained on Exc neurons achieved higher object discrimination accuracy and a closer alignment to the monkeys’ behavior, whereas decoders based on Inh neurons – though slightly less accurate – still carried significant category information and explained a unique fraction of the behavioral variance. Consistent with prior reports that excitatory neurons exhibit higher stimulus selectivity^8^, we confirm this classical finding in the inferior temporal cortex. However, we extend this observation by characterizing in greater depth the structure of response variability in each population. Specifically, we quantified response variance at multiple levels: image-by-image variability, variability within object categories, and variability across object categories. Interestingly, despite exhibiting higher stimulus and category selectivity, excitatory neurons showed lower response variance per image and reduced dispersion both within and across object categories compared to inhibitory neurons.

At first glance, this reduced variance might be expected to constrain representational richness. However, despite this lower variability, excitatory populations consistently achieved superior object decoding performance. Critically, this decoding advantage persisted even after explicitly controlling for differences in stimulus selectivity between the two populations. Moreover, excitatory-based decoders showed higher behavioral consistency, and this effect remained even when decoding accuracy was matched between populations. These findings suggest that the advantage of excitatory neurons cannot be reduced simply to higher selectivity or higher overall decoding accuracy.

To reconcile this apparent paradox — lower variance yet higher decodability and behavioral alignment — we examined the geometry of object representations using manifold analyses^49,50^. We found that excitatory populations exhibited smaller manifold radii and greater separation between object manifolds compared to inhibitory populations. In the framework of linear separability theory, smaller manifold radii combined with sufficient inter-manifold separation yield higher manifold capacity, enabling more object categories to be linearly decoded with fewer neurons. Thus, excitatory neurons appear to organize object representations in a more compact and geometrically favorable manner for downstream linear readout. In contrast, inhibitory populations displayed larger manifold radii and greater dispersion, which may reduce linear decodability while still contributing complementary, behaviorally relevant information.

It is still unclear how these distinct sources of information are readout and conveyed to the downstream brain areas like the PFC^55^, caudate^56^ and amygdala^57^. The exact nature of those readouts and therefore the individual contributions of the Exc and Inh neurons might be specific to the behaviors under study (e.g., object detection, facial emotion discrimination, or decision-making tasks such as go/no-go paradigms). Therefore, our study provides a template for generating linking hypotheses to test how sensory representation across cell types relate to behavioral tasks. Given the temporal variation in the amount of object information in the Exc and Inh neurons (inset Figure 5C), we speculate that a dynamic integration based decoding scheme that integrates information over time might be necessary to fully explain the object recognition behavior. Such temporally evolving integration mechanisms may differentially weight the geometrically compact representations carried by excitatory neurons and the more distributed representations carried by inhibitory neurons, thereby shaping the final behavioral output.

### Incorporating the role of Exc/Inh contributions in ANN-based brain models

While it remains common to seek an intuitive understanding of what image based features are differentially processed by these individual cell types, such inferences are often limited by the low-dimensional, apriori hypotheses (like size of receptive fields^58^, stimulus parameter tuning^9^) constructed by the experimenters. Complementary to this approach, our data provide two types of image-level constraints that might more effectively guide the next generation of brain models. These constraints are 1) reliably estimated image-by-image neural response of individual Exc and Inh neurons to a wide range of images 2) image-level differences in population decodes constructed from Exc and Inh neural activity. Access to these additional, finer grained, unbiased measurements could serve as target objectives in finetuning the ANN neural representations. Importantly, our results suggest that incorporating cell-type–specific constraints into ANN models should go beyond enforcing Dale’s principle alone. In addition to sign-constrained connectivity, excitatory and inhibitory populations in our data differ systematically in firing rate, response latency, reliability, variance structure, inter-neuronal correlations, decoding performance, and alignment with behavior. Artificial neural network units showed stronger alignment with excitatory neurons in representational similarity (CKA) as well as forward and reverse predictivity analyses, suggesting that current feedforward architectures may already approximate excitatory-dominated representations. However, these models typically lack explicit mechanisms that capture the complementary contribution of inhibitory populations to behavioral variance.

These observations imply that biologically grounded ANN models should incorporate structured inhibitory interactions that regulate variability, correlation structure, and trial-by-trial fluctuations, rather than treating inhibition as a purely global normalization mechanism. Explicitly modeling excitatory units as providing sparse, decorrelated, and linearly readable object representations, alongside inhibitory units that shape competition and behavioral variability, may improve both neural alignment and behavioral predictivity. In practice, this could involve constraining correlation structure, variance profiles, or temporal dynamics within model layers to better match cell-type–specific statistics observed in cortex.

A recent promising line of work^59^ has incorporated Dale’s principle (i.e., presynaptic neurons can only project as excitatory or inhibitory inputs on their postsynaptic partners) in a standard ANN architecture. Our results reported here serve as objective benchmarks for evaluating such models. But as mentioned earlier, this data can also be used during model development to regularize the neural networks, as demonstrated in previous work^60^.

### Implications for theories of Exc/Inh imbalance in various neurological disorders

A change in the Exc/Inh ratio is often deemed as an important neural marker for many neurological disorder ^61,62^. Similarly atypical sensory processing has also been a hallmark of many neurological disorders like autism spectrum disorder^63^, ADHD^64^, etc. Taken together, we speculate that understanding how participation of the Exc and Inh neurons shape the behavior of neurotypical adults, might be key in developing more robust theories of the differences that lead to clinical symptoms^65^. Importantly, our results highlight that Exc/Inh imbalance should be considered not only in terms of relative abundance or mean activity levels, but also in terms of how each population contributes to representational geometry, decoding efficiency, and behavioral predictivity. This perspective may help bridge the gap between cellular-level hypotheses and systems-level behavioral phenotypes. Selective targeting of Exc or Inh neurons with genetic techniques might help to model these disorders in experimental settings, uncovering novel therapeutic strategies. By causally manipulating one population at a time and quantifying changes in decoding accuracy, behavioral consistency, and image-specific error structure, future studies could determine whether clinical-like phenotypes arise from weakened excitatory readout, excessive inhibitory variability, or disrupted coordination between the two. Such experiments would move beyond global imbalance frameworks toward circuit-level explanations of sensory dysfunction. In this regard, it is important to also expand our study design to incorporate more specific cell types^66^ beyond the broad classifications of Inh and Exc neurons. Different subclasses of inhibitory interneurons, for example, may differentially influence correlation structure, timing, or gain control, and could therefore contribute distinctively to clinical phenotypes. A more fine-grained cell-type–specific approach may ultimately be necessary to link molecular and genetic perturbations to systems-level perceptual and behavioral outcomes.

### Future directions: Selective perturbation of excitatory and inhibitory neurons to test circuit level hypotheses

Our results allow us to generate testable predictions for future direct brain perturbation experiments that could selectively modulate Exc and Inh neurons. Recent advances in optogenetic^67^ and chemogenetic^68^ strategies allow to selectively transfect excitatory and inhibitory neurons and alter their responses. Such methods will allow us to explicitly test whether selective perturbation of Exc or Inh neurons can produce network changes like the balance of excitation and inhibition and reveal insights into brain disorders (as mentioned above). Specifically, if excitatory neurons form the principal substrate for downstream object readout, then selective suppression of excitatory neurons during behavior should produce a marked reduction in object discrimination accuracy and behavioral consistency. Because excitatory neurons exhibit lower correlation structure and reduced variance, perturbing them should degrade the reliability of the population code in a way that cannot be fully compensated by inhibitory activity.

In contrast, selective perturbation of inhibitory neurons may not simply reduce overall decoding performance, but instead alter response correlations, increase variability, or reshape the pattern of image-specific behavioral errors. Given that inhibitory populations explain a unique fraction of behavioral variance, their manipulation should modify behavioral signatures in ways that cannot be reproduced by uniformly scaling excitatory responses alone. Demonstrating such dissociable effects would provide causal evidence that excitatory and inhibitory neurons contribute distinct computational roles rather than differing only in degree of selectivity.

Understanding how Exc and Inh neurons contribute to the overall network activity will enable researchers to create detailed models of how neural circuits function. This can lead to a deeper comprehension of complex behaviors and cognitive processes.

## Materials and Methods

### Visual Stimuli

All stimuli used in this study were previously used in the Kar et al. 2019^16^ study and can be accessed at https://github.com/kohitij-kar/image_metrics. For a brief description of the stimuli, please refer below.

#### Generation of synthetic (“naturalistic”) images

High-quality images of single objects were generated using free ray-tracing software (http://www.povray.org), similar to Majaj et al. ^2^. Each image consisted of a 2D projection of a 3D model (purchased from Dosch Design and TurboSquid) added to a random background. The eight objects chosen were **bear**, **elephant**, **face**, **apple**, **car**, **dog**, **chair**, and **plane**. By varying six viewing parameters, namely position (*x* and *y*), rotation (*x*, *y*, and *z*), and size, we explored three types of identity-preserving object variation. For each object, we generated 80 images. Thus, we generated a total of 640 images. All images were achromatic with a native resolution of 256 × 256 pixels.

#### Subjects

Two adult male rhesus macaques (Macaca mulatta) participated in our experiments (monkeys N and B; age ∼10 years). All data were collected, and surgical and animal procedures were performed, in accordance with NIH guidelines, the Massachusetts Institute of Technology Committee on Animal Care, and the guidelines of Canadian Council on Animal Care on the use of laboratory animals and were also approved by the York University Animal Care Committee.

### Primate behavioral testing

#### Active binary object discrimination task

We measured monkey behavior from 2 male rhesus macaques. Images were presented on a 24-inch LCD monitor (1920 × 1080 at 60 Hz) positioned 42.5 cm in front of the animal. Monkeys were head fixed. Monkeys fixated on a white dot (0.2^°^) for 300ms to initiate a trial. The trial started with the presentation of a sample image (from a set of 640 images) for 100ms. This was followed by a blank gray screen for 100ms, after which the choice screen was shown containing a standard image of the target object (the correct choice) and a standard image of the distractor object. The monkey was allowed to freely view the choice objects for up to 1500ms and indicated its final choice by holding fixation over the selected object for 400ms. Trials were aborted if gaze was not held within ±2° of the central fixation dot during any point until the choice screen was shown. Prior to testing in the laboratory, monkeys were trained in their home-cages^11,69^ to perform the delayed match to sample tasks on the same object categories (but with a different set of images).

#### Passive Fixation Task

During the passive viewing task, monkeys fixated a white dot (0.2^°^) for 300ms to initiate a trial. We then presented a sequence of 5 to 10 images, each ON for 100ms followed by a 100ms gray (background) blank screen. This was followed by fluid (water) reward and an inter trial interval of 500ms, followed by the next sequence. All neural data analyzed for this study was collected when the animals (n=2) were performing the passive fixation tasks.

##### Eye Tracking

We monitored eye movements using video eye tracking (SR Research EyeLink 1000). Using operant conditioning and water reward, our 2 subjects were trained to fixate a central white square (0.2°) within a square fixation window that ranged from ±2°. At the start of each behavioral session, monkeys performed an eye-tracking calibration task by making a saccade to a range of spatial targets and maintaining fixation for 500ms. Calibration was repeated if drift was noticed over the course of the session.

Real-time eye-tracking was employed to ensure that eye jitter did not exceed ±2°, otherwise the trial was aborted, and data discarded. Stimulus display and reward control were managed using the MWorks Software (https://mworks.github.io).

#### Electrophysiological recording and data preprocessing

Prior to any neural recordings, we surgically implanted each monkey with a titanium headpost under aseptic conditions. We then chronically implanted 96-electrode Utah arrays (Blackrock Microsystems) in the IT cortex of the two monkeys: three in the right cerebral hemisphere of monkey B, and two in the right hemisphere of monkey N. Each array consisted of a 4.4mm x 4.2mm square grid of electrodes spaced 400um apart. Each electrode (silicon shanks) in the array was 1.5mm long, with platinum tips. The arrays were pneumatically inserted and placed inferior to the superior temporal sulcus and anterior to the posterior middle temporal sulcus. We have previously analyzed multi-unit activity from these same monkeys with a significantly higher yield of reliable neural sites^16,17,60^. However, to obtain well-isolated single units we made conservative choices (see below), for this study, that restricted our analyses to a total of 199 single neurons (monkey N: 64, monkey B: 135) pooled across the two monkeys.

#### Sorting multi-unit neurons to single neurons

During each experimental session, multi-unit neural activity was recorded continuously at a sampling rate of 20 kHz using an Intan RHD Recording Controller (Intan Technologies, LLC). Offline, all processing was performed using the SpikeInterface (version 0.98.2) framework. The raw voltage signals were pre-processed by applying a bandpass filter (400 Hz to 5 kHz, fifth-order Butterworth filter), following which multi-unit spikes were detected using thresholding. A multi-unit spike event was defined as the threshold crossing when voltage (falling edge) deviated by more than five times the median absolute deviation of the raw voltage values. The multi-unit spikes were then sorted into single-units using Tridesclous (TDC; version 1.6.8), accessed via SpikeInterface. We picked Tridesclous as it had the lowest reported false positive error rate compared to other spike sorters (see Figure 2 in Buccino et al., 2020^70^). We obtained 277 single neurons from TDC spike sorter.

We then performed some automatic curation steps to ensure that the identified single neurons were well-isolated. For each of the single neurons, we removed duplicated spikes by iteratively computing the first spike and removing any spike less than 1ms apart from it. This step was done as a neuron is incapable of producing action potentials very close to each other due to its biophysical refractory period, and the biophysical refractory window is often assumed to be 1 ms^71^. We then computed three quality metrics for each of the single neurons: inter-spike interval (ISI) violations ratio, presence ratio, and amplitude cutoff. The ISI violations ratio^40^ measures the rate of refractory period violations within a specified window, and is computed as,

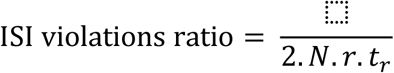

With 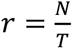 is the firing rate and where *N* the total number of spikes in the spike train, *T* is the total duration of the recording in seconds, *t*_*r*_ is the refractory period threshold (here set to 1.2ms) and *N*_*viol*_ is the number of inter-spike intervals shorter than *t*_*r*_. A higher number of ISI violations ratio represents greater contamination. For example, a value of 0.5 would indicate that 50% of the spikes of the neuron are contaminated. We only retained neurons with ISI violations ratio less than 0.4.

Then, we measure the presence ratio. It measures the fraction of time bins in which at least 1 spike occurred, and is computed as,

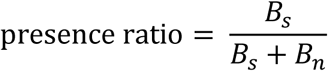

where *B*_*s*_ is the number of bins in which spikes occurred, and *B*_*n*_ is the number of bins in which no spikes occurred. Each bin is defined to be 60s long. This quantity ranges between 0 and 1, with low values indicating missing spikes or highly selective firing patterns. We only retained neurons with presence ratio greater than 0.9.

Finally, the amplitude cutoff measures the fraction of spikes that are likely to be missing from the cluster due to amplitude thresholding during spike detection. This metric estimates the proportion of the spike amplitude distribution that falls below the detection threshold. It is computed by first constructing the distribution of spike amplitudes for a given unit. Assuming the true amplitude distribution is approximately symmetric, the missing fraction is estimated as the area of the distribution that would lie below the detection threshold. Formally, the amplitude cutoff is defined as:

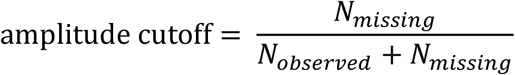

where *N*_observed_ is the number of detected spikes and *N*_missing_ is the estimated number of spikes falling below threshold, inferred from the truncated tail of the amplitude distribution. This quantity ranges between 0 and 1, with larger values indicating greater loss of low-amplitude spikes and therefore poorer cluster isolation. We retained only neurons with amplitude cutoff less than 0.1. Thus, using quality metrics with conservative thresholding, we retained 199 well-isolated single neurons out of 277 units.

##### Neuron cell-type classification based on spike waveform

For each neuron, we quantified two waveform features from its average action potential (AP), namely peak-to-valley and recovery time. Peak-to-valley time is defined as the time interval between the global negative trough of the waveform and the subsequent positive peak. Recovery time is the time interval between the positive peak and the return of the waveform to baseline. Several past studies have observed a bimodal distribution of peak-to-valley time, corresponding broadly to narrow-spiking and broad-spiking units, which are commonly interpreted as putative inhibitory and excitatory neurons^8,25,28,34,41^. Because waveform features such as peak-to-valley time and recovery time are strongly correlated with neuronal cell type, they have been widely used to classify extracellularly recorded units into putative excitatory and inhibitory populations ^8,25,34,35,37–39,41,66,72^. To determine the number of distinct waveform-based clusters in our dataset, we fit Gaussian mixture models (GMMs) with varying numbers of components and used the Bayesian Information Criterion (BIC) for model selection. The BIC identified a three-component model as the best fit to the joint distribution of peak-to-valley time and recovery time. We therefore fit a three-component GMM to these two features and assigned neurons to clusters based on maximum posterior probability. Based on waveform width, clusters with narrow peak-to-valley times were classified as putative inhibitory neurons, whereas clusters with broader waveforms were classified as putative excitatory neurons. This resulted in a dataset composed of 162 excitatory neurons (Monkey B: 121, Monkey N: 41) and 31 inhibitory neurons (Monkey B: 9, Monkey N: 22).

#### Data Analyses

##### Behavioral Metrics (B.I_1_)

We have used a one-vs-all image level behavioral performance metric (**B.I_1_** similar to the one used in ^11,16^) to quantify the behavioral performance of the monkeys (described below, see Figure SX). This metric estimates the overall discriminability of each image containing a specific target object from all other objects (pooling across all 7 possible distractor choices). Critically, our task structure yields, for each image, an image-specific decision (or confusion) pattern across distractor categories (here 7). For each trial of the task, a specific sample image was shown, and a binary choice task screen was presented. So, the data obtained from each trial is a correct (1) or incorrect (0) choice from the subjects. Each trial can be labeled with 2 unique identifiers – a unique sample image (one out of 640) and a unique task (one out of seven possible tasks given the image, e.g. bear vs. dog).

Given an image of object ‘***i***’, and all seven distractor objects (distractor ‘j’, *j* ≠ *i*) we computed the average performance per image (each element of the **B.I_1_** vector) as,

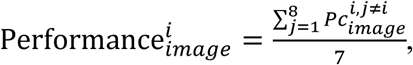

where *P_C_* refers to the fraction of correct responses for the binary task between objects ‘***i***’ and ‘*j*’. While the **B.I_1_** vector provides an estimate of monkeys’ image-by-image accuracy, the overall performance accuracy can be determined by taking an average of the **B.I_1_** vector (across all images).

To compute the reliability of this vector, we split the trials per image into two equal halves by resampling without substitution. The median of the Spearman-Brown corrected correlation of the two corresponding vectors (one from each split half), across 1000 repetitions of the resampling was then used as the reliability score (i.e., internal consistency).

#### Neural recording metrics per IT neuron

##### Baseline Firing Rates

Baseline firing rates were estimated by counting the number of spikes evoked by 26 gray (blank screen) images in the 70-170ms time period after Test image onset and multiplying by 100ms to obtain the number of spikes within 1 second.

##### Maximum Evoked Firing Rates

Maximum evoked firing rates per neuron were estimated by counting the number of spikes evoked by an image stimulus that produced the maximum response in the 70-170ms time period after Test image onset and multiplying by 100ms to obtain the number of spikes within 1 second.

##### Image rank-order response reliability per neuron

To estimate the reliability of the responses per neuron given a time window, we divided neural responses along the trial (repetition) dimension into two independent halves. For each split, neural responses were averaged across trials to obtain mean firing rates per neuron and image. Reliability is computed independently for every neuron (across all 640 images). Reliability is quantified as the Pearson correlation between the two splits. To correct for the use of half the available data in each split, raw split-half correlations (*r*) were adjusted using the Spearman-Brown^73^ formula:

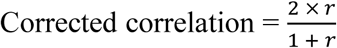

Split-half estimation was repeated multiple times with independent random splits, and the mean corrected correlation across repetitions was used as the final reliability estimate.

##### Latency

Response latency was defined independently for each neuron based on the temporal evolution of its image rank-order reliability. For each time bin following stimulus onset, split-half reliability was computed as described above (Spearman–Brown corrected Pearson correlation across the 640 images). Latency was defined as the earliest time point after stimulus onset at which the neuron’s split-half reliability exceeded a threshold (0.2 in the main text, see **Figure S2** for other threshold values). This criterion identifies the first moment at which the neuron exhibits a stable and reproducible image-selective response across repetitions.

##### Response Variance across images

Response variance was estimated per neuron as the variance in the spike count across 640 images in the 70-170ms time period after Test image onset.

##### Stimulus Selectivity

Stimulus selectivity was quantified for each neuron using multiple complementary measures commonly used in visual neurophysiology. All measures were computed across the full set of 640 images using trial-averaged firing rates within the 70-170ms time window.

Let *R*_*i*_ denote the mean firing rate of a neuron to stimulus *i*, with *n* total stimuli. For each neuron, the following measures were computed:

- **Depth of Selectivity (DOS)**^45,46^

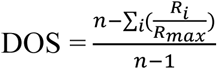 where *R*_*max*_ is the maximum firing rate across stimuli. DOS ranges from 0 to 1, with higher values indicating stronger selectivity (i.e., responses concentrated on fewer stimuli). Neurons with negligible dynamic range (i.e., *R*_*max*_ ≤ 0) were assigned a DOS of 0. We reported this quantity in Figure 2 but computed additional statistics on the following measures.
- **Breadth of Selectivity**^47^ Responses were first normalized for each neuron 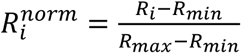, where *R*_*min*_ and *R*_*max*_ are the minimum and maximum responses across stimuli. Breadth was then defined as:

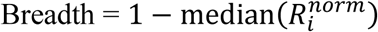 This metric ranges from 0 to 1, with higher values indicating greater selectivity. Neurons with no response variability (i.e., *R*_max_ = *R*_min_) were assigned a breadth of 0.
- **Selectivity index (SI)**^48^

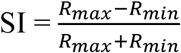 where *R*_*max*_ and *R*_*min*_ are the maximum and minimum firing rates across stimuli. Under non-negative firing rates, SI ranges from 0 to 1, with larger values indicating stronger stimulus selectivity. If the denominator was zero, SI was set to 0.

##### Category Selectivity

Category selectivity^55^ quantified the extent to which a neuron responded more differently to stimuli from different object categories than to stimuli within the same category. For each neuron, we computed:

- **Between-category difference (BCD):** the average absolute difference in firing rate between all pairs of stimuli belonging to different object categories.
- **Within-category difference (WCD):** the average absolute difference in firing rate between all pairs of stimuli belonging to the same category.

Category selectivity was then defined as:

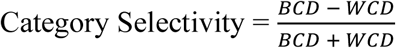

This metric ranges from −1 to 1. Positive values indicate greater differentiation between categories than within categories, consistent with category-selective responses. Values near zero indicate no category structure in the response profile. We compared the category selectivity of neurons from each population across different time windows.

#### Population-level analyses

##### Reliability-matched sampling

Excitatory and inhibitory neuronal populations differed both in the number of recorded units and in their response reliability. Because reliability can directly influence decoding performance and representational metrics, we constructed reliability-matched populations prior to all population-level analyses. For each comparison, neurons were subsampled such that excitatory and inhibitory populations had matched reliability distributions. Reliability was defined per neuron as the Spearman–Brown corrected split-half correlation (see above). Neurons were first restricted to the overlapping range of reliabilities shared by the two populations. We then performed stratified sampling across reliability quantiles to ensure comparable coverage of the reliability spectrum in both groups while maintaining equal population sizes. All population-level analyses were repeated across independent random draws of matched populations, and reported results reflect averages across these reliability-controlled subsamples.

##### Estimation of IT population decode accuracies (***N.I_1_***)

To estimate what information downstream neurons could easily “read” from a given IT neural population, we used simple, biologically plausible linear decoders (i.e., linear classifiers), that have been previously shown to link IT population activity and primate behavior^2^. Such decoders are simple in that they can perform binary classifications by computing weighted sums (each weight is analogous to the strength of synapse) of input features and separate the outputs based on a decision boundary (analogous to a neuron’s spiking threshold). Here we have used a linear discriminant analysis-based classifier. The learning model generates a decoder with a decision boundary that is optimized to best separate images of the target object from images of the distractor objects. Instead of directly minimizing a regularized loss function, Linear Discriminant Analysis (LDA) estimates class-specific means and a shared covariance matrix across classes and computes a linear decision boundary that maximizes the ratio of between-class variance to within-class variance. We trained 8 one-vs-all classifiers (one per object), each distinguishing one target object from all other objects. After training each of these classifiers with a 10-fold cross validation, we generated a class probability score (*sc*) per classifier for all held out test images. The train and test sets were pseudo-randomly chosen multiple times until every image of our image set was part of the held-out test set. We then computed the binary task performances, by calculating the percent correct score for each pair of possible binary tasks given an image. For instance, if an image was from object *i* (Target), then the percent correct score for the binary task between object *i* and object *j* (Distractor), *Pr*^*i*,*j*^ was computed as,

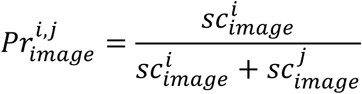

From each percent correct score, we then estimated a neural **N.I_1_** score (per image), following the same procedures as the behavioral metric.

##### Estimation of **N.I_1_** across varying population size

To evaluate how neural image-level performance (**N.I_1_**) scales with population size, we computed **N.I_1_** separately for excitatory and inhibitory populations while systematically varying the number of neurons included in the decoder. For each population size *n*, we constructed reliability-matched subsets of excitatory and inhibitory neurons (see *Reliability-matched sampling*). Reliability matching was performed independently for each population size to ensure that comparisons between excitatory and inhibitory populations were not confounded by differences in reliability distributions. For each sampled population, firing rates were trial-averaged within the analysis window and used to train linear decoders (see section above). From these decoders, we computed both the neural image-level behavioral metric (**N.I_1_**). This procedure was repeated across multiple independent random samplings (40 repetitions per population size) to account for variability due to subsampling. Reported results reflect averages across these repetitions.

##### Estimation of **N.I_1_** across varying Exc/Inh Ratios

To assess how population composition influences neural image-level performance (**N.I_1_**), we constructed neural populations with fixed total size while systematically varying the excitatory/inhibitory (Exc/Inh) ratio. Because the inhibitory pool contained 31 neurons, total population size was fixed at 31 units across all conditions to ensure that differences in performance reflected population composition rather than neuron count.

Population construction involved two stages of sampling. First, to control for reliability differences between excitatory and inhibitory neurons, we generated reliability-matched pools of excitatory and inhibitory units (see *Reliability-matched sampling*). This step was repeated across independent random draws (n=20) to account for variability in matched subsets.

Second, for each reliability-matched pool, we generated populations with varying Exc/Inh ratios. We began with fully inhibitory populations and progressively replaced inhibitory neurons with excitatory neurons, increasing the number of excitatory units from 0 to 31 while correspondingly decreasing inhibitory units. For each ratio, neurons were randomly sampled without replacement from the matched excitatory and inhibitory pools (n=20 repetitions).

For every Exc/Inh ratio, the **N.I_1_** metric was computed on each sampled population. This two-level resampling procedure (reliability matching followed by ratio-specific subsampling) was repeated multiple times (total = 400), and reported results reflect averages across repetitions.

##### Consistency of **N.I_1_**with **B.I_1_**

To quantify how well neural image-level performance (**N.I_1_**) captured behavioral image-level performance (**B.I_1_**), we computed a noise-corrected correlation (“consistency”) between neural and behavioral image-level sensitivity profiles. For each analysis time window, we compared the excitatory and inhibitory **N.I_1_** with the measured behavioral performance (**B.I_1_**, see *Behavioral Metrics*). For each window and each population, we computed the Pearson correlation between the **B.I_1_** and **N.I_1_**. Because both neural and behavioral measurements are noisy, we applied noise-correction by dividing the raw correlation by the square root of the product of the neural reliability and behavioral reliability:

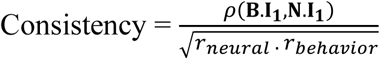

where *ρ* is the Pearson correlation across images, *r*_neural_ is the reliability of the neural **N.I_1_** estimate for that condition, and *r*_behavior_ is the reliability of the behavioral **B.I_1_** measurements (both computed via split-half methods; see *Behavioral Metrics* and *Neural Recording Metrics* sections). To quantify variability due to resampling, we also computed consistency separately for each repetition of the **N.I_1_** within a time window and reported dispersion using the median absolute deviation (MAD).

##### Partial correlation of **N.I_1_**with **B.I_1_**

To test whether the consistency between neural and behavioral image-level performance was specific to each cell type (as opposed to reflecting shared variance between excitatory and inhibitory populations), we computed partial correlations between behavioral performance (**B.I_1_**) and neural performance (**N.I_1_**) while controlling for the other population.

To estimate both reliability and cross-population dependence robustly, we used a split-half procedure on neural trials. For each time window, we first constructed reliability-matched excitatory and inhibitory populations (30 neurons per population). For each matched draw, neural trials were randomly shuffled and split into two independent halves, and trial-averaged firing rates were computed separately for each split. We then computed **N.I_1_** independently for each split, yielding two independent neural **N.I_1_** estimates per population (e.g., 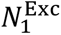 and 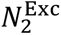, and similarly for inhibitory).

We then computed two types of partial correlations across images. First, we computed a within-population “floor” estimate by correlating behavior with one neural split while controlling for the other split of the same population:

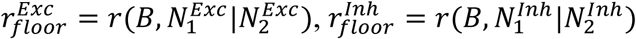

This floor provides a conservative baseline driven by measurement noise and split-half variability. Second, we computed cross-population partial correlations to assess the unique behavioral variance explained by each population after accounting for the other population:

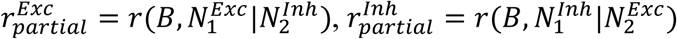

All partial correlations were computed across images using Pearson partial correlation. The overall procedure was repeated across multiple independent draws of reliability-matched populations (n=10) and trial splits (n=10), producing distributions of floor and partial-correlation estimates for each time window.

##### Object manifold geometry metrics

To characterize the representational geometry of excitatory and inhibitory populations, we analyzed object manifolds constructed from trial-averaged population responses within a fixed post-stimulus time window. For each object (8 objects; 80 images per object), we defined a manifold as the set of neural response vectors to all images belonging to that object. Each manifold was therefore represented as a matrix *X* ∈ ℝ^*M*×*N*^, where *M* = 80 images and *N* denotes the number of neurons in the reliability-matched population.

Prior to geometric analysis, neural responses were z-scored per neuron across images and globally mean-centered to remove baseline offsets. To ensure fair comparisons between excitatory and inhibitory populations, all manifold metrics were computed on reliability-matched subsamples of equal size (n=30, see Reliability-Matched Sampling). The full procedure was repeated across bootstrap samples to quantify variability.

For each object manifold, we computed the following geometric quantities:

- **Manifold center and Radius** The manifold center was defined as the mean response vector across images, 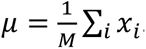. The absolute radius was defined as the root-mean-square distance of samples to the center:

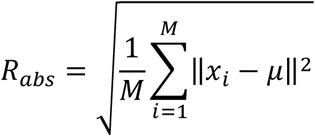 We additionally computed the center norm ‖*μ*‖ and defined the relative radius as:

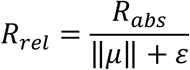 where *ε* is a small constant for numerical stability. The relative radius captures the spread of the manifold relative to its mean magnitude.
- **Effective Dimensionality** Manifold dimensionality was quantified using the participation ratio (PR) of the covariance matrix:

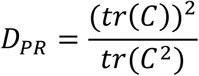 where *C* is the covariance of the manifold responses. The participation ratio provides an estimate of the effective number of dimensions spanned by the manifold.
- **Manifold Capacity** We estimated manifold classification capacity as a deterministic function of relative radius and participation-ratio dimensionality:

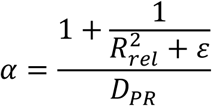 Under this formulation, capacity increases when manifolds are compact relative to their center magnitude (small *R*_rel_) and decreases as manifolds occupy higher-dimensional subspaces (large *D*_PR_). Capacity was computed independently for each object manifold and then averaged across the 8 objects to yield a population-level estimate.

In addition to within-manifold geometry, we quantified the distance between manifolds by computing the mean pairwise Euclidean distance between manifold centers across objects. We further derived a scale-invariant separation index by normalizing center separation by the mean manifold radius.

All geometric metrics (relative radius, participation ratio, capacity, and separation measures) were computed separately for excitatory and inhibitory populations within each bootstrap iteration and subsequently averaged across bootstrap samples.

##### Across and within object variance

To quantify how strongly neurons differentiate objects relative to variability across exemplars, we computed across-object and within-object variance for each neuron separately.

Neural responses were first trial-averaged within the analysis time window, yielding a response matrix with 640 images (8 objects × 80 images) by neurons. All variance measures were computed independently for each neuron.

**Across object variance** was defined as the variance across object means. For each neuron, we first computed its mean response to each of the 8 objects (averaging across the 80 images per object). We then computed the sample variance of these 8 object means. This measure captures how strongly the neuron’s average response differs across objects.

**Within object variance** was defined as the average variance across exemplars within each object. For each neuron and each object, we computed the sample variance of responses across the 80 images belonging to that object. We then averaged these variances across the 8 objects. This measure captures how variable the neuron’s responses are across different exemplars of the same object.

All analyses were performed separately for excitatory and inhibitory populations.

##### Inter-neuron correlations

To quantify shared response structure within each population, we computed inter-neuron correlations across images separately for excitatory and inhibitory neurons. Neural responses were first averaged within the analysis time window and then trial-averaged, yielding for each population an image-by-neuron response matrix (640 images × neurons).

For each population, we computed the Pearson correlation between the response vectors of every pair of distinct neurons across the 640 images. This produced a neuron-by-neuron correlation matrix (excluding the diagonal). For each neuron, we summarized its average inter-neuron correlation by taking the mean correlation with all other neurons.

#### Predictivity analyses

##### Model selection

We selected a broad and diverse set of 38 artificial neural network (ANN) models spanning a wide range of architectures, training objectives, and inductive biases. These models include classic convolutional networks such as AlexNet^53^, VGG16^74^, and multiple ResNet^54^ variants (e.g., ResNet-18, ResNet-50, ResNet-101, ResNet-152), as well as deeper architectures like ConvNeXt^75^, DenseNets^76^, Inception-v1/v3^77,78^, and NASNet/PNASNet^79,80^. We also included lightweight models such as MobileNet^81^, ShuffleNet^82^, and SqueezeNet^83^, biologically inspired recurrent models like CORnet-S and CORnet-RT^84^, the intermediate architectures from the ConvRNN suite^85^ and modern transformer-based architectures including Vision Transformers^86^ (ViT) and Swin Transformers^87^. To capture the effects of training, we incorporated models trained under standard supervised learning, self-supervised learning (e.g., SimCLR^88^, SwAV^89^), and robust models developed by adversarial training^90^. All models were pre-trained on the ImageNet^91^ object classification dataset and task (except for minor differences in training specifics as per the original publications), and we used the publicly available pre-trained weights.

##### Layer selection

For each ANN, we extracted activations from a single high-level layer corresponding to the macaque IT cortex, as defined by the ANN-IT mapping on Brain-Score^14^. The layer selection in Brain-Score is done by asking which layer of the model best predicts existing macaque IT data, using data from a previous publication^2^, and methods previously published^22^. The IT layer of recurrent models (CORNet and ConvRNNs) was directly labeled by the authors of the respective papers (“IT” in CORNet^84^ and “conv8” for ConvRNNs^85^) This layer produces an M-dimensional feature vector per image, where M varies by model (e.g., 4096 for AlexNet, 100372 for ResNet-50). For a stimulus set of N images, this yielded an M × N matrix of model responses per ANN (See **<u>Figure</u> SX** for a full list of models and their IT layers). Of note, we did not use the same neural data used in layer selection for our analyses in the article. Neural data used in this article was collected in a separate set of monkeys using a different image-set.

All predictivity analyses were performed using the predictivity toolbox^92,93†^

##### Prediction of neural responses from ANN features

We modeled each IT neural site as a linear combination of the ANN model features. Using a 50%/50% train/test split of the images, we then estimated the regression weights (i.e., how we can linearly combine the model features to predict the neural site’s responses) using a ridge regression procedure. The neural responses used for training (R^TRAIN^) and testing (R^TEST^) the encoding models were averaged firing rates (measured at the specific sites) within the time window (70 to 170ms in Figure 7, 180 to 240ms in **Figure SX**) post stimulus onset. The training images used for regressing the model features onto a neuron, at each time point, were sampled randomly (repeats included random subsampling) from the entire image set. For each set of regression weights (*w*) estimated on the training image responses (R^TRAIN^), we generated the output of that ‘synthetic neuron’ for the held-out test set (M^PRED^) as,

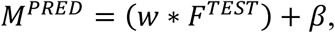

where *w* and *β* are estimated via the PLS regression and *F*^*TEST*^ are the model activation features for the test image-set.

The percentage of explained variance, *IT predictivity* for that neural site, was then computed by normalizing the correlation between the prediction and the neural response for that site by the self-consistency of the test image responses 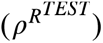 for that site and the self-consistency of the regression model predictions 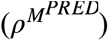 for that site (estimated by a Spearman Brown corrected trial-split correlation score).

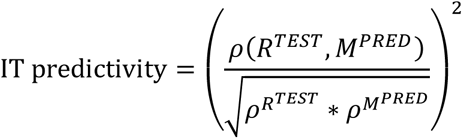

##### Prediction of ANN features from neural responses

To assess the extent to which neural responses predict ANN feature representations, we performed the reverse analysis. In this case, ANN feature activations were modeled as linear combinations of neural responses using the same 50%/50% train/test splitting procedure and ridge regression.

The regression weights were estimated on the training images, mapping neural responses to ANN features. Predicted model features for the held-out test images were then generated using the learned weights. Performance was again quantified as noise-corrected explained variance between predicted and actual ANN features.

Because ANN activations are deterministic (i.e., there is no trial-to-trial variability), reliability correction in the reverse analysis only accounts for noise in the neural measurements. Thus, prediction reliability reflects variability introduced by neural sampling and regression estimation.

##### CKA between ANN features and neural responses

To compare representational similarity between ANN feature spaces and IT population responses, we used centered kernel alignment^94^ (CKA). Neural responses were computed as trial-averaged firing rates within the analysis window (70 to 170ms in Figure 7, 180 to 240ms in **Figure SX**). For each population (excitatory and inhibitory), we formed an image-by-neuron response matrix by averaging across trials, yielding one response vector per image and neuron.

For each model, we computed CKA between the model feature matrix and the neural response matrix separately for excitatory and inhibitory populations, producing one CKA value per model and population (CKA_Exc_, CKA_Inh_). To estimate the reliability (noise ceiling) of CKA within each neural population, we performed a split-half procedure across trials: trials were randomly permuted and divided into two halves, responses were averaged within each half to form two independent image-by-neuron matrices, and CKA was computed between these two matrices. This yielded a split-half CKA reliability estimate for excitatory and inhibitory populations (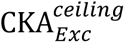, 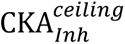). The final noise-corrected CKA per population was computed as 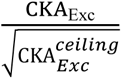 and 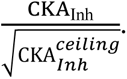

#### Statistical Tests to determine significance of effects reported

##### Statistical Testing

All statistical comparisons were guided by a consistent decision procedure based on the distributional properties and pairing of the data. For parametric test, we first assessed whether each distribution was approximately normal using the Shapiro-Wilk test^95^. If both groups satisfied the normality assumption, we applied parametric tests. For paired data (e.g., Exc vs. Inh predictions across models), we used a paired t-test (two-tailed; degrees of freedom = n – 1). For unpaired comparisons (e.g., between Excitatory and Inhibitory populations), we used an independent samples t-test (degrees of freedom = n₁ + n₂ – 2).

If at least one distribution violated the normality assumption, we used non-parametric alternatives. Specifically, we applied the Wilcoxon signed-rank test for paired comparisons and the Wilcoxon rank-sum test (equivalent to the Mann–Whitney U test) for unpaired data.

For non-parametric tests, we employed permutation tests, where a null distribution was first obtained by randomly shuffling the labels of the two variables under comparison.

All tests were two-tailed unless stated otherwise. We report exact p-values and test statistics throughout.

## Acknowledgments

KK was supported by the Canada Research Chair Program (CRC-2021-00326) and the Simons Foundation Autism Research Initiative (SFARI, 967073)-Pilot award, NSERC Discovery Program (NSERC, RGPIN-2024-06223), and funds from the Azrieli Foundation and Brain Canada Foundation (2023-0259). SM was supported by Connected Minds (CFREF funded program at York University) Postdoctoral fellowship.

## Supplementary Material

**Figure S1.**
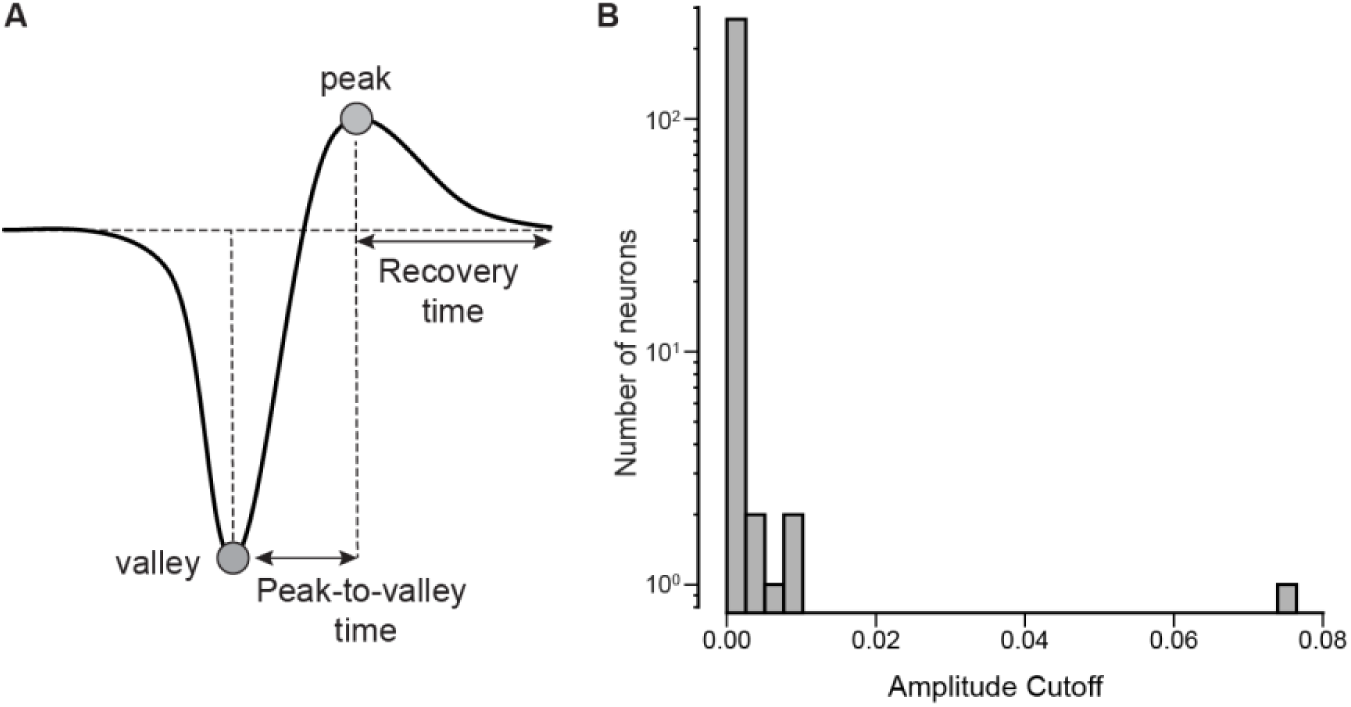
Spike waveform features and amplitude cutoff distribution. (**A**) Schematic of the average action potential waveform illustrating the two features used for cell classification. Peak-to-valley time was defined as the interval between the global minimum (valley) and the subsequent positive maximum (peak). Recovery time was defined as the interval from the positive peak to the return to baseline. (**B**) Distribution of amplitude cutoff values across recorded neurons. All units exhibited low amplitude cutoff values (<0.1), indicating minimal truncation of spike waveforms and supporting the quality of single-unit isolation.

**Figure S2.**
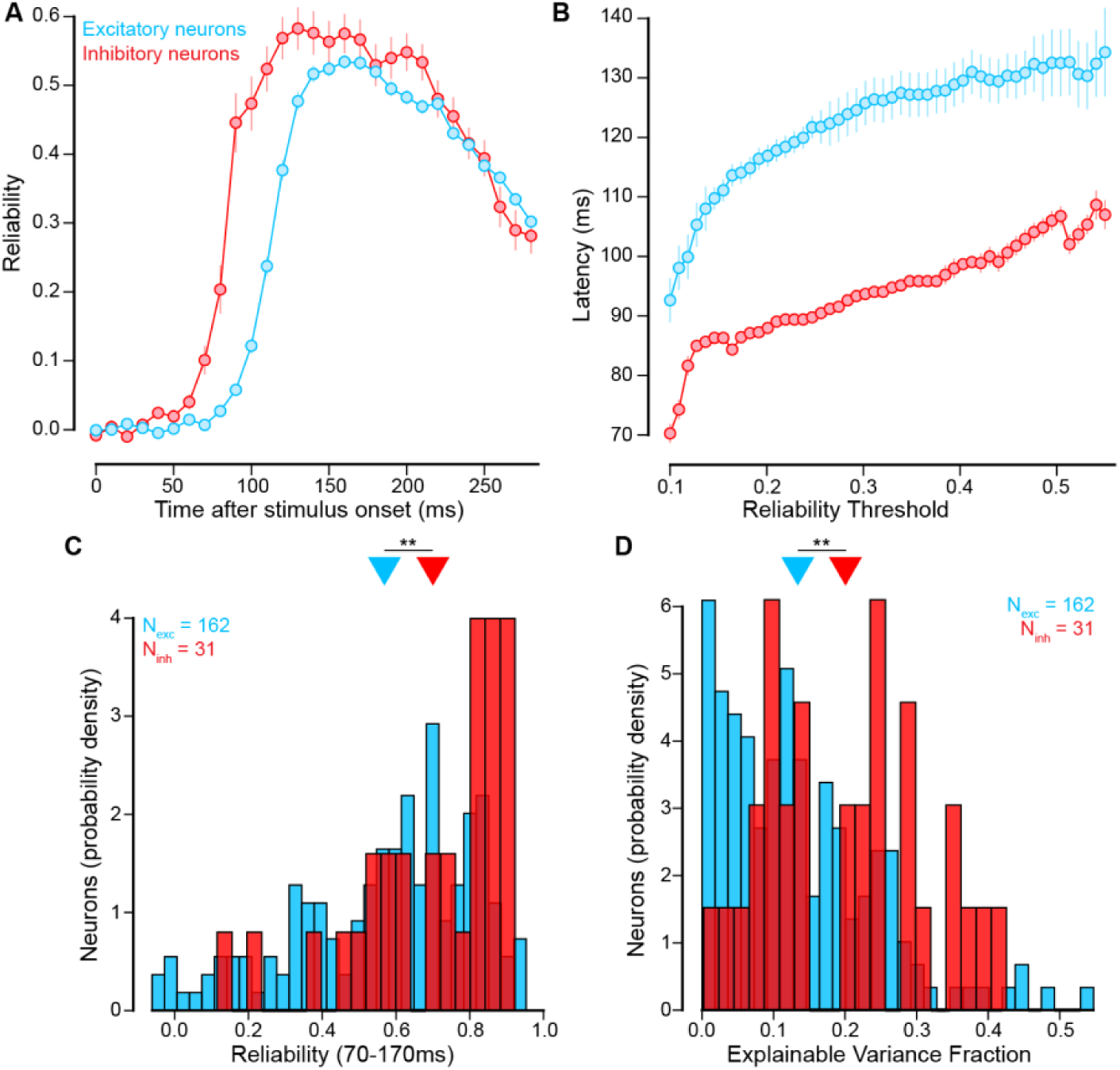
Reliability, response latency, and explainable variance of excitatory and inhibitory neurons. **(A)** Time course of split-half reliability following stimulus onset for excitatory (blue) and inhibitory (red) neurons. Reliability rises earlier for inhibitory neurons, but reaches higher peak values for excitatory neurons. **(B)** Response latency as a function of reliability threshold. Excitatory neurons always exhibit longer response latencies than inhibitory neurons. (A-B) Points denote mean ± MAD across neurons (n=162 Exc neurons, n=31 Inh neurons). **(C)** Distribution of neuronal reliability computed in the 70–170 ms post-stimulus window. Excitatory neurons (n = 162) exhibited significantly lower reliability (mean=0.57) than inhibitory neurons (n = 31, mean=0.70, two-sided unpaired Wilcoxon test across neurons: z=3.032, p=0.002). **(D)** Distribution of explainable variance fraction (EVF). Excitatory neurons showed significantly lower EVF (mean=0.13) compared to inhibitory neurons (mean=0.20, two-sided unpaired Wilcoxon test across neurons: z=3.106, p=0.002. **(B-D)** **p < 0.01.

**Figure S3.**
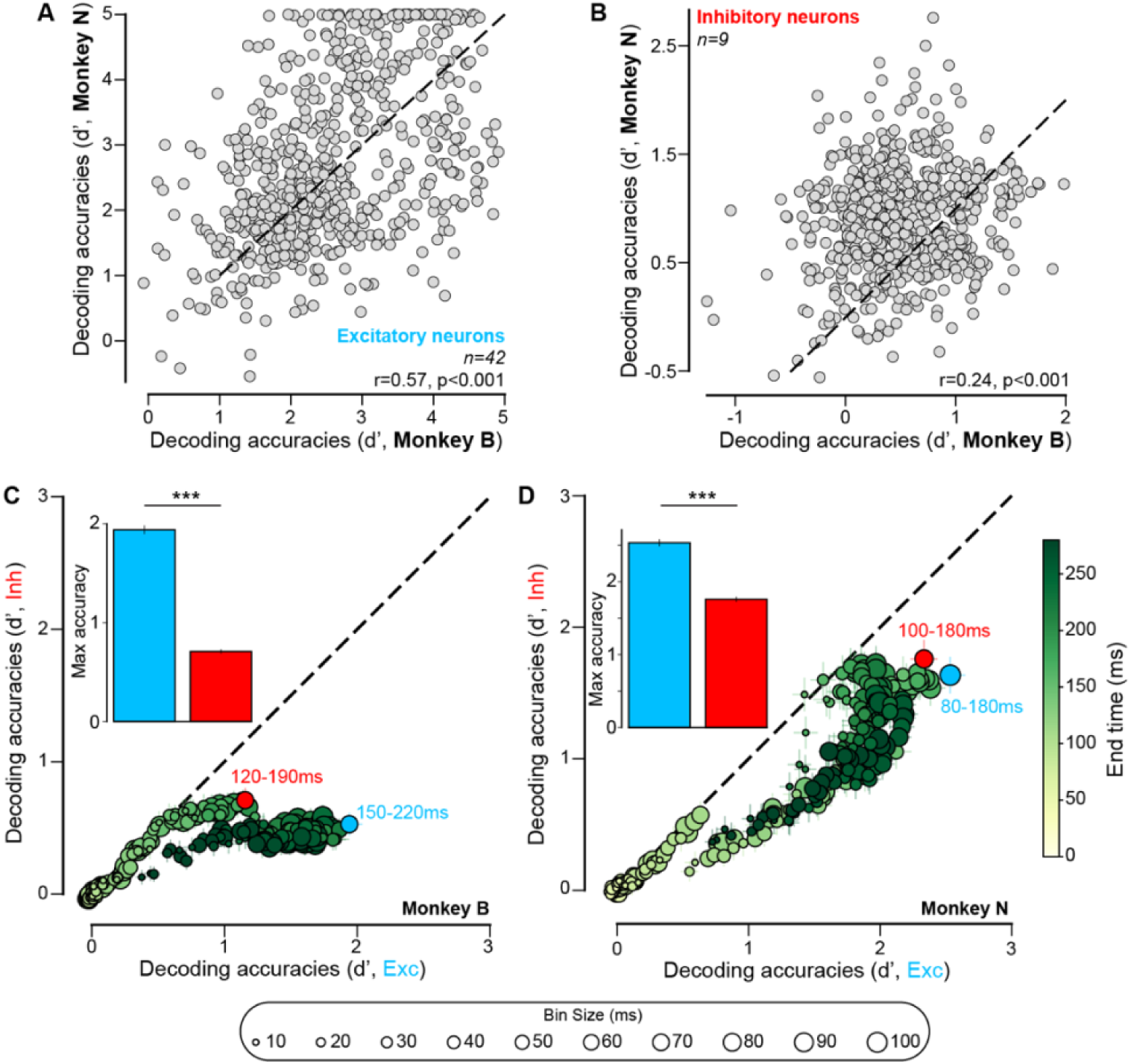
Comparison of decoding performance between excitatory and inhibitory neuronal populations across monkeys. (**A**) Cross-monkey consistency of decoding accuracy for excitatory neurons. Each point represents the decoding accuracy (d′) obtained from the 70-170ms time window for Monkey B (x-axis) and Monkey N (y-axis). To enable a fair comparison across monkeys, the excitatory population in Monkey B was randomly subsampled multiple times (n=20) to match the number of excitatory neurons recorded in Monkey N (n = 42). Excitatory population decoders showed strong consistency across monkeys (noise-corrected Pearson correlation: *r(638) = 0.57, p < 0.001*; n = 640 images). (**B**) Same analysis as in (A) for inhibitory neurons. The inhibitory population in Monkey N was randomly subsampled repeatedly (n=20) to match the number of inhibitory neurons recorded in Monkey B (n = 9). Decoding accuracies showed weaker but significant cross-monkey consistency (noise-corrected Pearson correlation: *r(638) = 0.24, p < 0.001*; n = 640 images). (**C-D**) Same as **Figure 3C** for individual monkeys. (**C**) Comparison of decoding performance between excitatory (x-axis) and inhibitory (y-axis) populations for Monkey B across temporal integration windows. Each point represents a decoder defined by a specific bin size (circle size) and window end time (color scale, see **Figure 3A**). Colored points highlight the best-performing integration windows for each population (Exc: 150-220ms, Inh: 120-190ms). The inset shows the maximum decoding accuracy achieved by each population, revealing significantly higher peak performance for excitatory neurons (two-sided paired Wilcoxon test across images: *z=2432.0, p<0.001*, n=640 images). (**D**) Same analysis as in (**C**) for Monkey N, showing consistently higher decoding performance for excitatory compared to inhibitory populations across integration windows. Insets again report the maximum decoding accuracy for each population (Exc: 80-180ms, Inh: 100-180ms, two-sided paired Wilcoxon test across images: *z=26162.0, p<0.001*, n=640 images) (**C-D**) ***p < 0.001.

**Figure S4.**
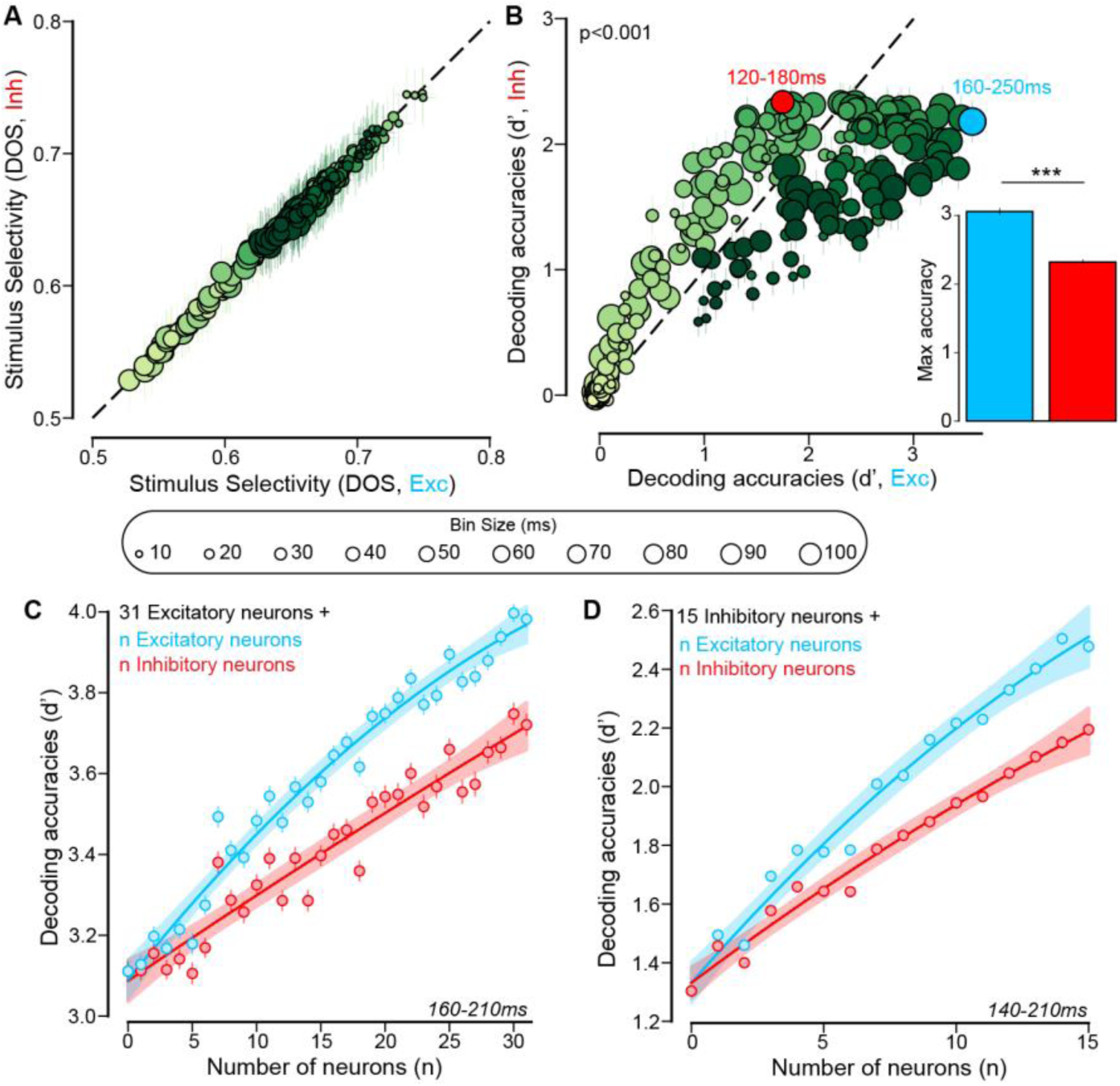
Control analyses on factors influencing decoding accuracy. (**A**) Stimulus-selectivity matching. For each temporal integration window (circle size indicates bin size; color indicates window end time), we subsampled excitatory neurons such that the excitatory population’s stimulus selectivity (DOS) matched that of the inhibitory population for the same window. Points show the resulting DOS values for the matched excitatory (x-axis) and inhibitory (y-axis) populations. (**B**) Decoding after selectivity matching. Using the selectivity-matched excitatory subsamples from (**A**), we compared object decoding accuracy (d′) between excitatory (x-axis) and inhibitory (y-axis) populations across the same temporal integration windows. Despite matching DOS, excitatory populations achieved higher decoding accuracy overall (two-sided paired Wilcoxon test across conditions: *z=11077.0, p<0.001*, n=245 conditions). The inset summarizes the maximum decoding accuracy across windows for each population (two-sided paired Wilcoxon test across images: *z=37687.0, p<0.001*, n=640 images, ***p < 0.001). Colored markers denote the best-performing windows for each population (Exc: 120-180ms, Inh: 160-250ms). (**C**) Decoding accuracy (mean ± MAD across images) as a function of population size when combining a reliability-matched subset of excitatory neurons (n = 31) with an increasing number of inhibitory neurons (red) or excitatory neurons (blue) at the 160-120ms time window (best Exc accuracy). Adding Exc neurons led to a greater improvement in decoding performance compared to adding inhibitory neurons (at n=31, two-sided paired Wilcoxon test across images: *z=47686.0, p<0.001*, n=640 images). (**D**) Same analysis as in (**C**), but starting from a subset of inhibitory neurons (n = 15) and progressively adding excitatory or inhibitory neurons. Performance improved more when adding excitatory neurons than when adding inhibitory neurons (at n=31, two-sided paired Wilcoxon test across images: *z=38955.0, p<0.001*, n=640 images).

**Figure S5.**
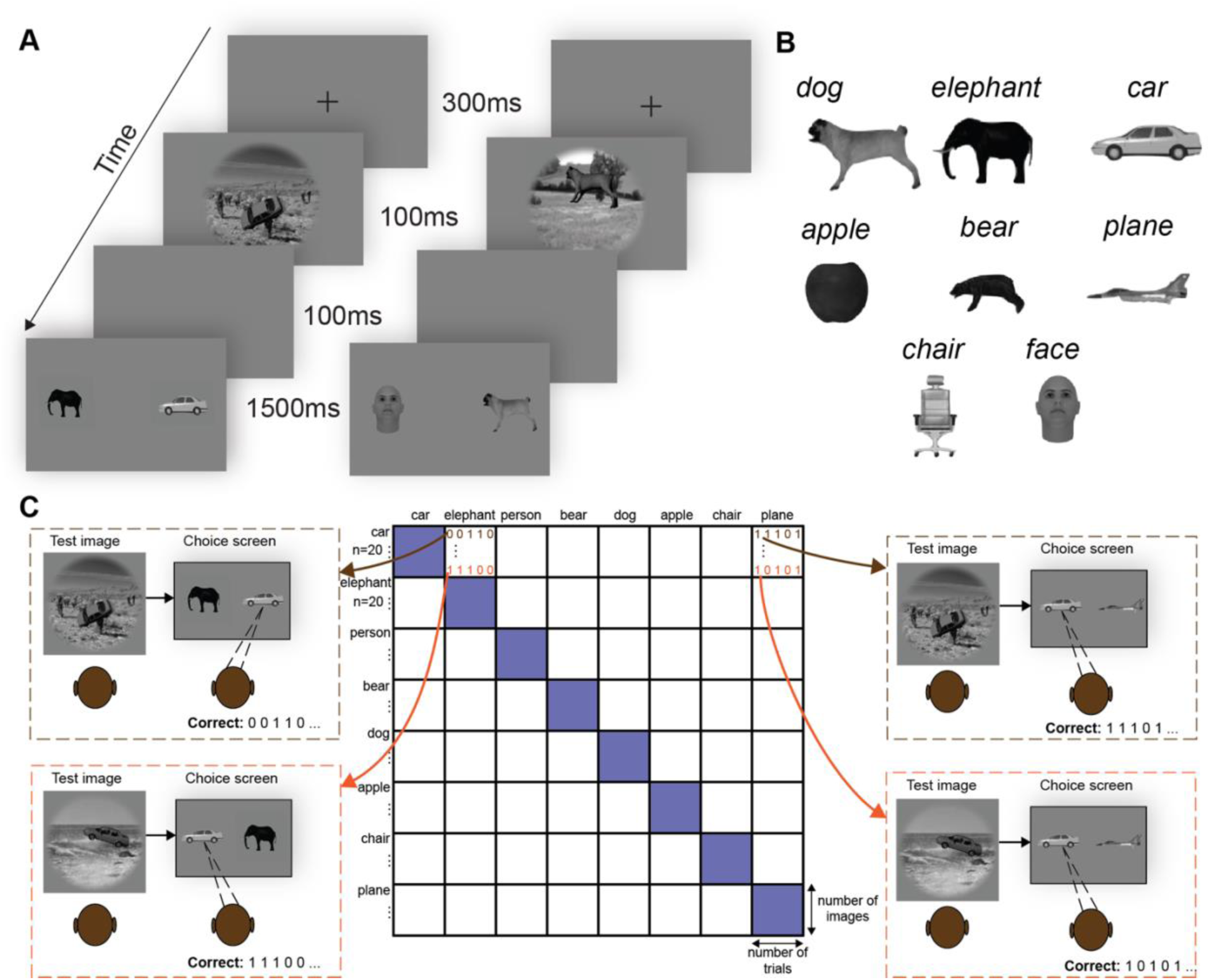
Behavioral task design and computation of image-level behavioral metrics. (**A**) Trial structure of the two-alternative forced-choice (2AFC) object discrimination task. Each trial began with a fixation period (300ms), followed by presentation of the test image (100ms). After a short delay (100ms), a choice screen appeared for 1500ms showing two object choices, one target and one distractor. Subjects have to indicated which object was present in the image. (**B**) Object categories used in the experiment. Eight objects were tested: dog, elephant, car, apple, bear, plane, chair, and face. (**C**) Illustration of how behavioral responses were aggregated to compute image-level performance metrics. For each test image, performance was measured across multiple binary discrimination tasks against different distractor objects. The resulting response patterns were arranged into an object-by-object matrix where rows correspond to target objects and columns correspond to distractor objects. Each cell summarizes behavioral performance across images and trials for that pairwise discrimination, which together define the image-level behavioral signature used in subsequent analyses.

**Figure S6.**
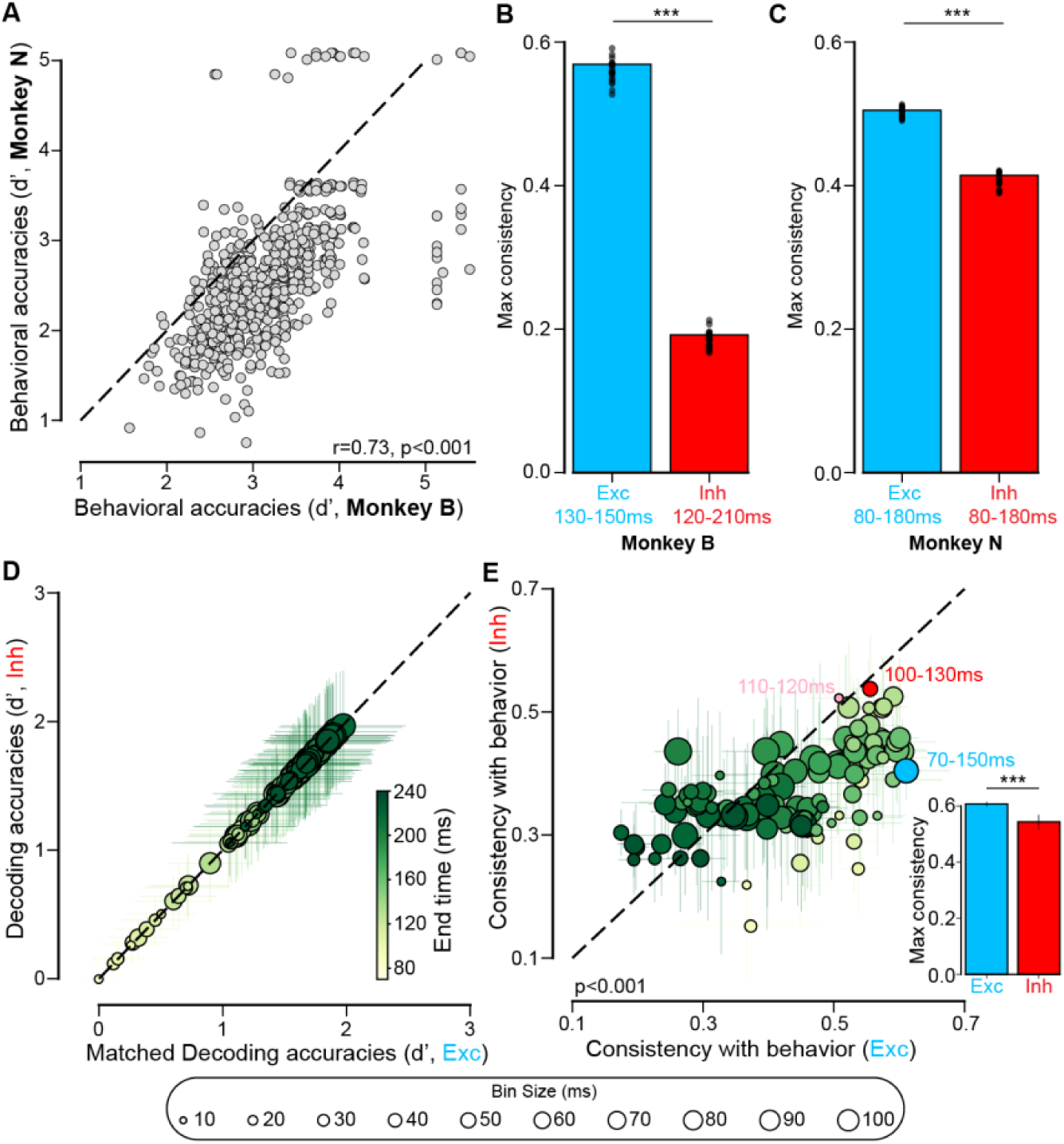
Relationship between neural decoding performance and behavioral accuracy across monkeys. (**A**) Cross-monkey consistency of behavioral performance. Each point represents the behavioral accuracy (d′) for a given image (n=640 images) measured in Monkey B (x-axis) and Monkey N (y-axis). Behavioral performance was significantly correlated across monkeys (noise-corrected Spearman correlation: *r(638) = 0.73, p < 0.001*). (**B**) Maximum consistency between neural decoding accuracy and behavioral performance for Monkey B. Bars show the peak correlation obtained across temporal integration windows for excitatory (blue) and inhibitory (red) populations. The highest consistency for excitatory neurons occurred at 130–150ms, whereas inhibitory neurons peaked at 120–210ms (permutation test across, n=20, reliability-matched samplings: *D=0.374*, *p<0.001*). (**C**) Same analysis as in (**B**) for Monkey N. Excitatory decoders achieved higher behavioral consistency than inhibitory decoders (permutation test across, n=20, reliability-matched samplings: *D=0.095*, *p<0.001*). The best excitatory window and best inhibitory window both occurred at 80–180ms. (**D**) Control analysis using accuracy-matched samples of excitatory neurons. For each temporal window, excitatory neurons were sampled (n=31) such that their decoding accuracy matched that of inhibitory decoders. Each point represents one such matched decoder (mean ± MAD across samplings). (**E**) Behavioral consistency of Exc **N.I_1_** (x-axis) vs. Inh **N.I_1_** (y-axis) given accuracy-matched sampled Exc populations. Each point corresponds to a decoding condition defined by a temporal integration window (mean consistency ± MAD across samplings). Colors indicate window end time and circle size indicates bin size. Despite matching accuracy, excitatory populations achieved higher consistency with behavior overall (two-sided paired Wilcoxon test across conditions: *z=1113.0, p<0.001*, n=245 conditions). The inset shows the maximum behavioral consistency achieved by excitatory and inhibitory decoders. Excitatory decoders reached higher peak consistency with behavior than inhibitory decoders (permutation test across samplings: *D=0.072, p<0.001*, n=20 samplings). (**B-E**) ***p < 0.001.

**Figure S7.**
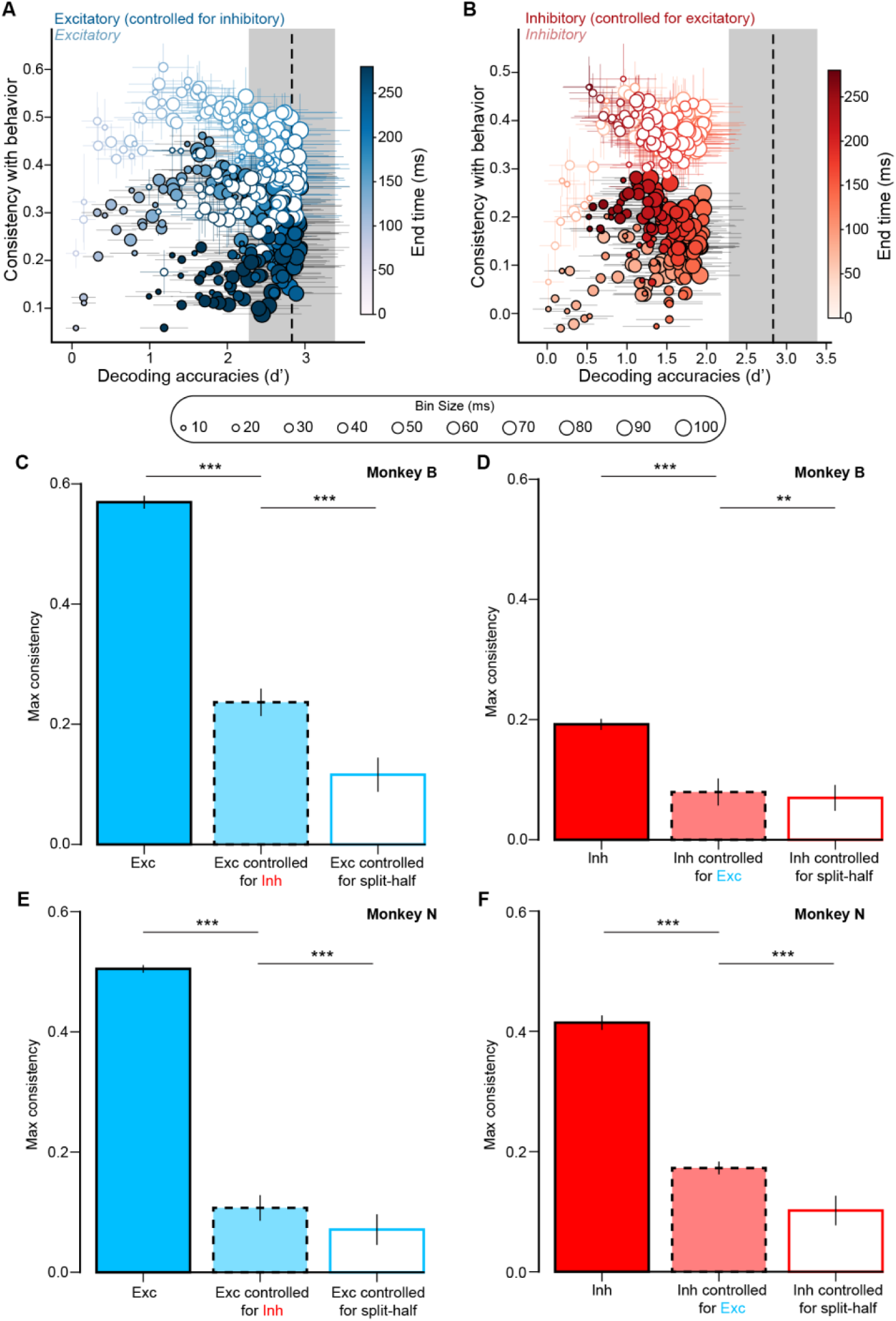
Partial correlation as a function of decoding accuracy and across individual monkeys. (**A**) Relationship between decoding accuracy (x-axis) and behavioral consistency for excitatory decoders. Whited-filled circles show the correlation between excitatory **N.I_1_** and behavior, while color-filled circles show the partial correlation after controlling for inhibitory **N.I_1_**. Each point represents a decoder defined by a temporal integration window (bin size shown by circle size; end time shown by color). The dashed vertical line indicates the behavioral accuracy of the monkeys. (**B**) Same analysis as in (**A**) but for inhibitory decoders. White-filled circles show the correlation between inhibitory decoder predictions and behavior, while color-filled circles show the partial correlation after controlling for excitatory decoder predictions. (**C**) Maximum behavioral consistency of Exc **N.I_1_** for Monkey B across conditions. Bars show the maximum consistency achieved by excitatory decoders (Exc), excitatory decoders controlled for inhibitory predictions (Exc controlled for Inh; partial correlation), and a split-half control within the excitatory population. Partial correlations remained significantly higher than the split-half control, indicating that excitatory decoders explain unique variance in behavior beyond inhibitory decoders (Exc>Exc controlled for Inh: *D=0.323, p<0.001*, Exc controlled for Inh > Exc controlled for split-half: *D=0.120, p<0.001*). (**D**) Same analysis as in (**C**) for Inh decoders (Inh>Inh controlled for Exc: *D=0.106, p<0.001*, Inh controlled for Exc > Inh controlled for split-half: *D=0.01, p=0.004*). (**E**) Maximum behavioral consistency for Exc **N.I_1_** in Monkey N. Bars show excitatory decoders (Exc), excitatory decoders controlling for excitatory predictions (Exc controlled for Inh), and a split-half control within the excitatory population. Partial correlations remained significantly higher than the split-half control, indicating that excitatory decoders also explain unique variance in behavior beyond inhibitory decoders (Exc>Exc controlled for Inh: *D=0.395, p<0.001*, Exc controlled for Inh > Exc controlled for split-half: *D=0.036, p<0.001*). (F) Same analysis as in (**E**) for Inh decoders (Inh>Inh controlled for Exc: *D=0.234, p<0.001*, Inh controlled for Exc > Inh controlled for split-half: *D=0.071, p<0.001*). (**C-F**) Two-sided permutation test across samplings (n=20), ***p < 0.001, **p < 0.01.

**Figure S8.**
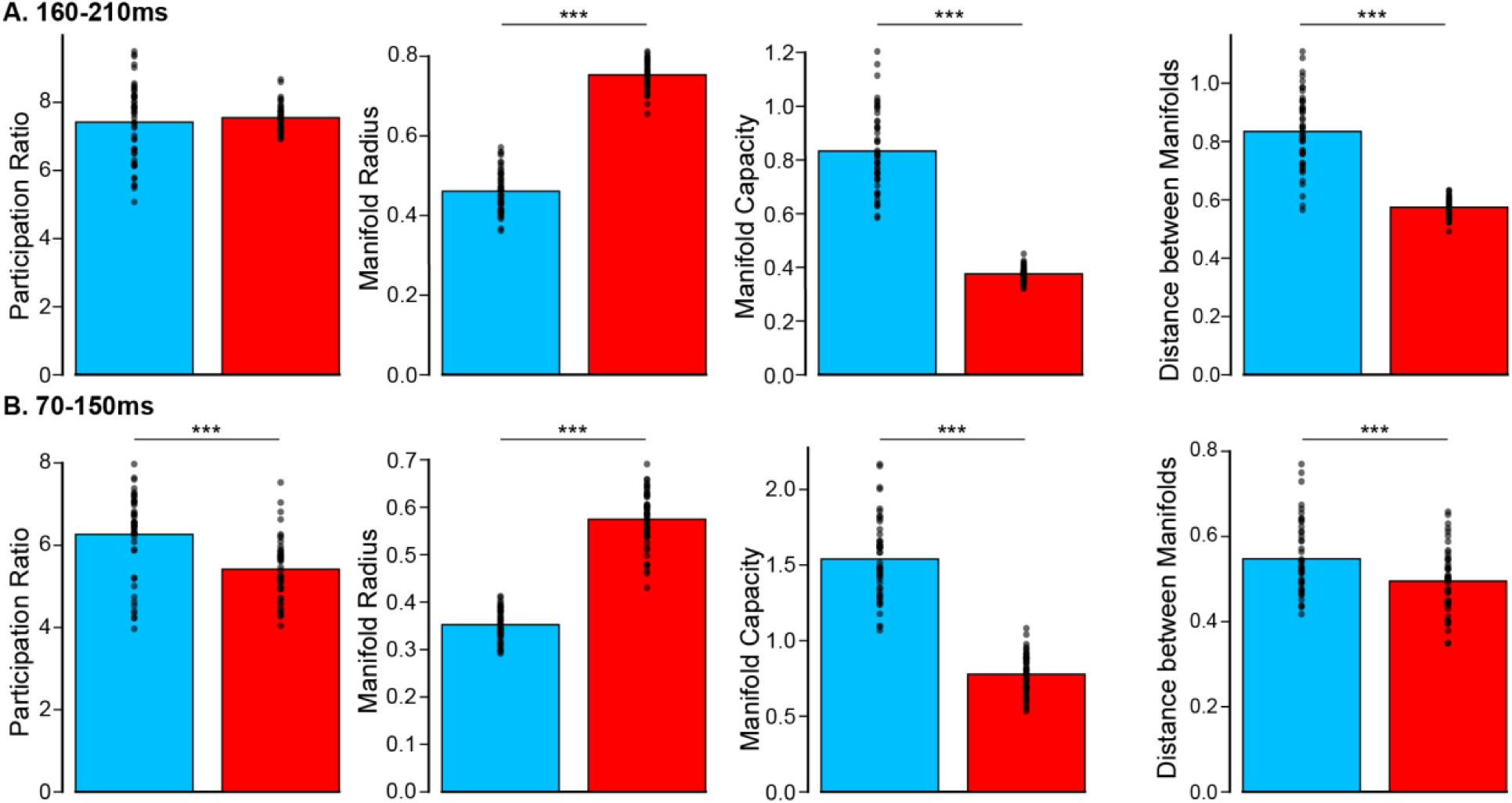
Comparison of manifold geometry between excitatory and inhibitory populations across time windows. (**A**) Manifold geometry metrics computed in the 160–210 ms time window (time window leading to the best decoding accuracy). From left to right: participation ratio (effective dimensionality), manifold radius, manifold capacity, and distance between manifolds for excitatory (blue) and inhibitory (red) populations. Bars indicate the mean across multiple reliability-matched samplings (n=50); Points represent individual samplings. In this time window, inhibitory populations exhibited larger manifold radius but lower manifold capacity and inter-manifold distance compared to excitatory populations (permutation tests across samplings; participation ratio: *D=-0.133, p=0.272*, radius: *D=-0.293, p < 0.001*, capacity: *D=0.457, p < 0.001*, distance: *D=0.260, p < 0.001*, ***p < 0.001). (**B**) Same analysis as in (**A**), computed in the 70–150 ms time window (time window leading to the best consistency with behavior). Excitatory populations showed higher effective dimensionality, larger manifold capacity, and greater inter-manifold distance, whereas inhibitory populations exhibited larger manifold radius (permutation tests across samplings; participation ratio: *D=0.846, p < 0.001*, radius: *D=-0.223, p < 0.001*, capacity: *D=0.762, p < 0.001*, distance: *D=0.053, p < 0.001*, ***p < 0.001).

**Figure S9.**
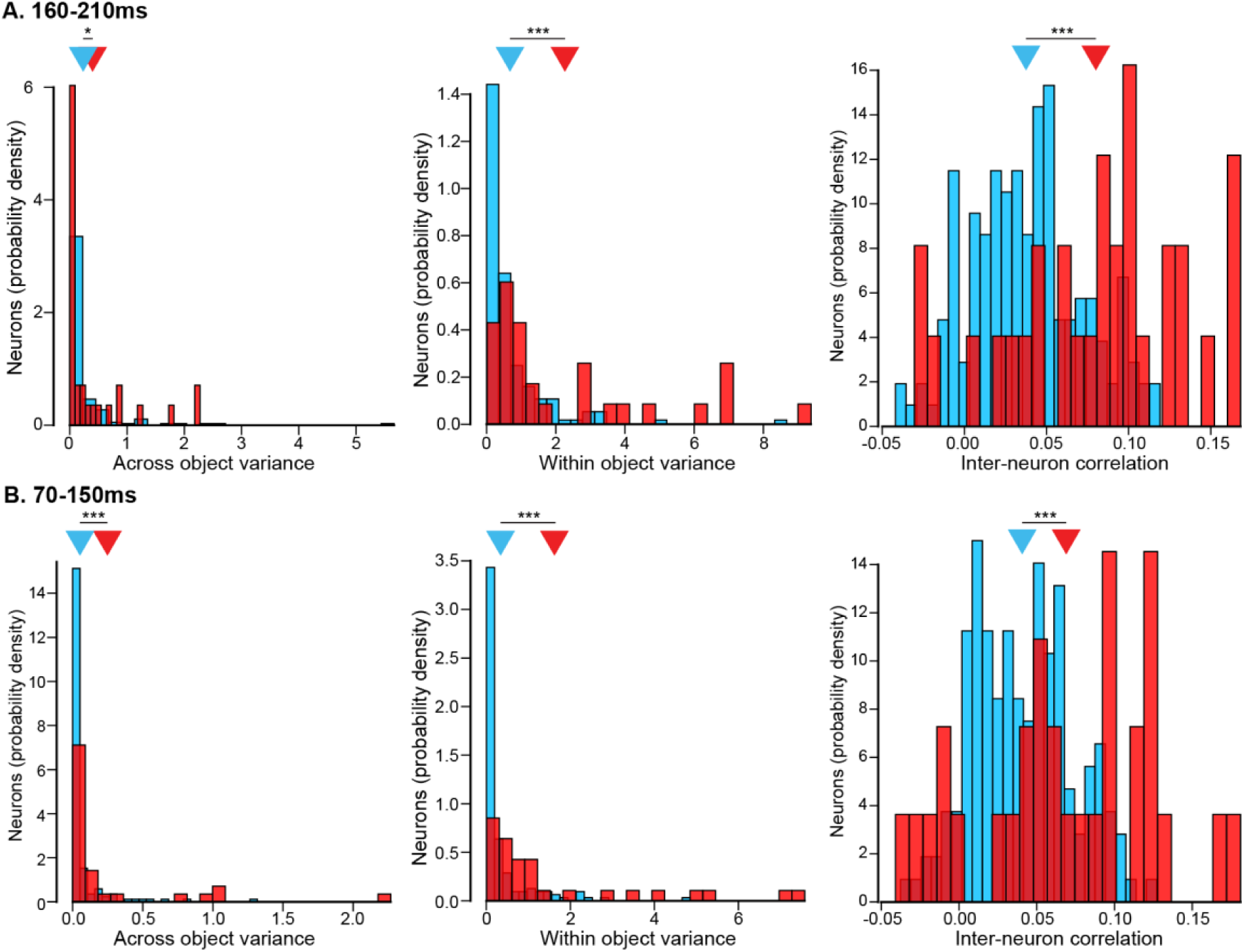
Variance structure and inter-neuronal correlations of excitatory and inhibitory populations. (**A**) Distribution of across-object variance (left), within-object variance (middle), and pairwise inter-neuron correlations (right) for excitatory (blue) and inhibitory (red) neurons in the 160–210ms time window (time window leading to the best decoding accuracy). Histograms show probability density across neurons. Inhibitory neurons exhibited significantly higher across (two-sided unpaired Wilcoxon test: *z=2.025, p=0.043*) and within-object variance (two-sided unpaired Wilcoxon test: *z=4.436, p < 0.001*) and stronger pairwise correlations (two-sided unpaired t-test: *t(191)=-5.714, p < 0.001)*. (**B**) Same analysis as in (**A**), computed in the 70–150 ms time window (time window leading to the best consistency with behavior). Inhibitory neurons exhibited significantly higher across (two-sided unpaired Wilcoxon test: *z=4.601, p < 0.001*) and within-object variance (two-sided unpaired Wilcoxon test: *z=5.159, p < 0.001*) and stronger pairwise correlations (two-sided unpaired t-test: *t(191)=-3.891, p < 0.001)*. (**A-B**) N = 162 excitatory neurons, N = 31 inhibitory neurons, *p < 0.05, ***p < 0.001.

**Figure S10.**
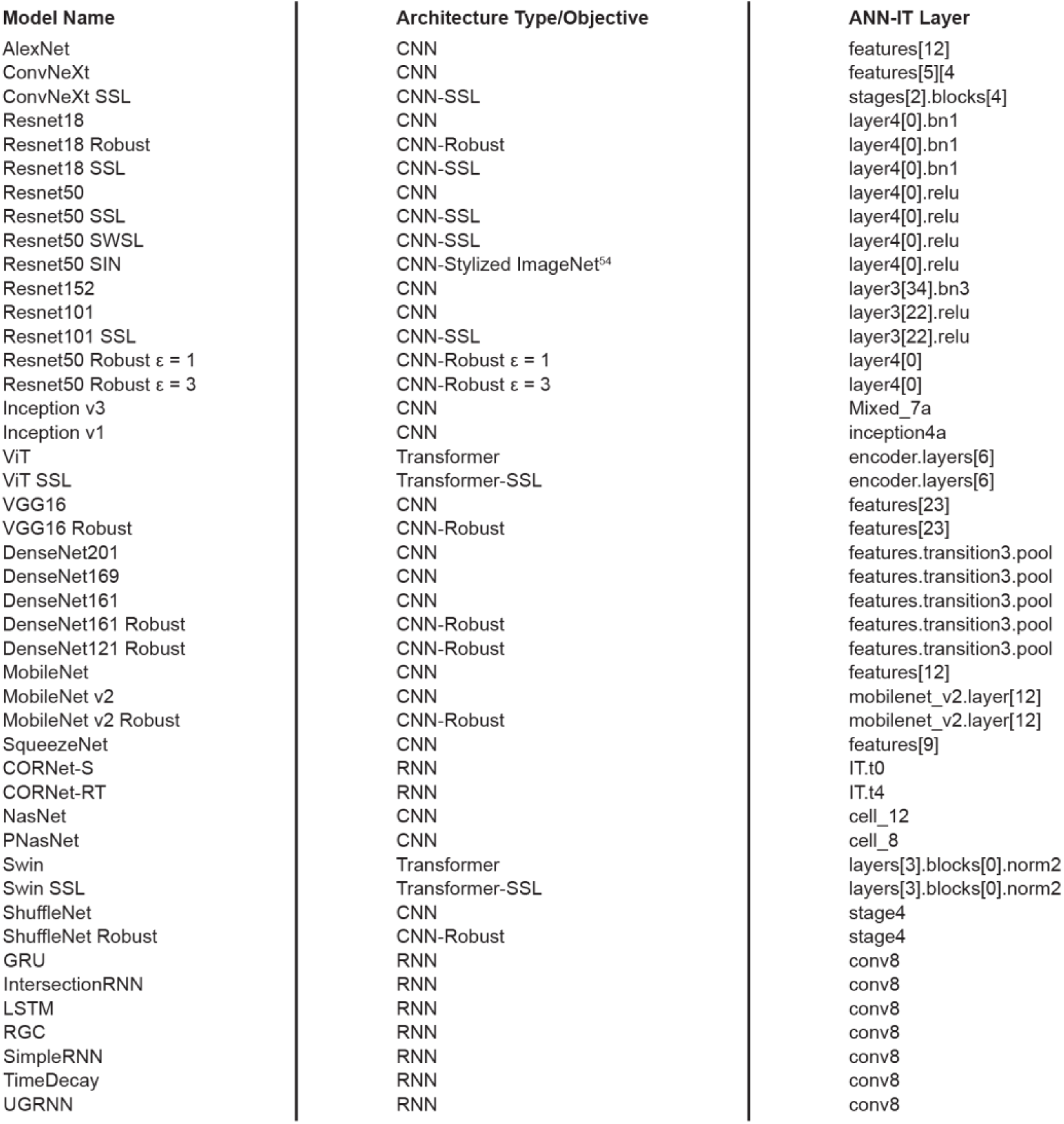
Artificial neural network models and corresponding IT-aligned layers used for comparison with neural data. Table summarizing the artificial neural network (ANN) models included in the analyses, their architecture type and training objective, and the specific layer used for comparison with IT cortex responses. Models span convolutional neural networks (CNNs), self-supervised variants (CNN-SSL, Transformer-SSL), robustness-trained models (CNN-Robust; varying adversarial strengths ε), recurrent neural networks (RNNs), and transformer-based architectures. For each model, we selected the layer that best corresponded to IT representations (ANN-IT layer), according to Brain-Score. This diverse model set enabled evaluation of how architectural class and training objective relate to alignment with IT neural responses.

**Figure S11.**
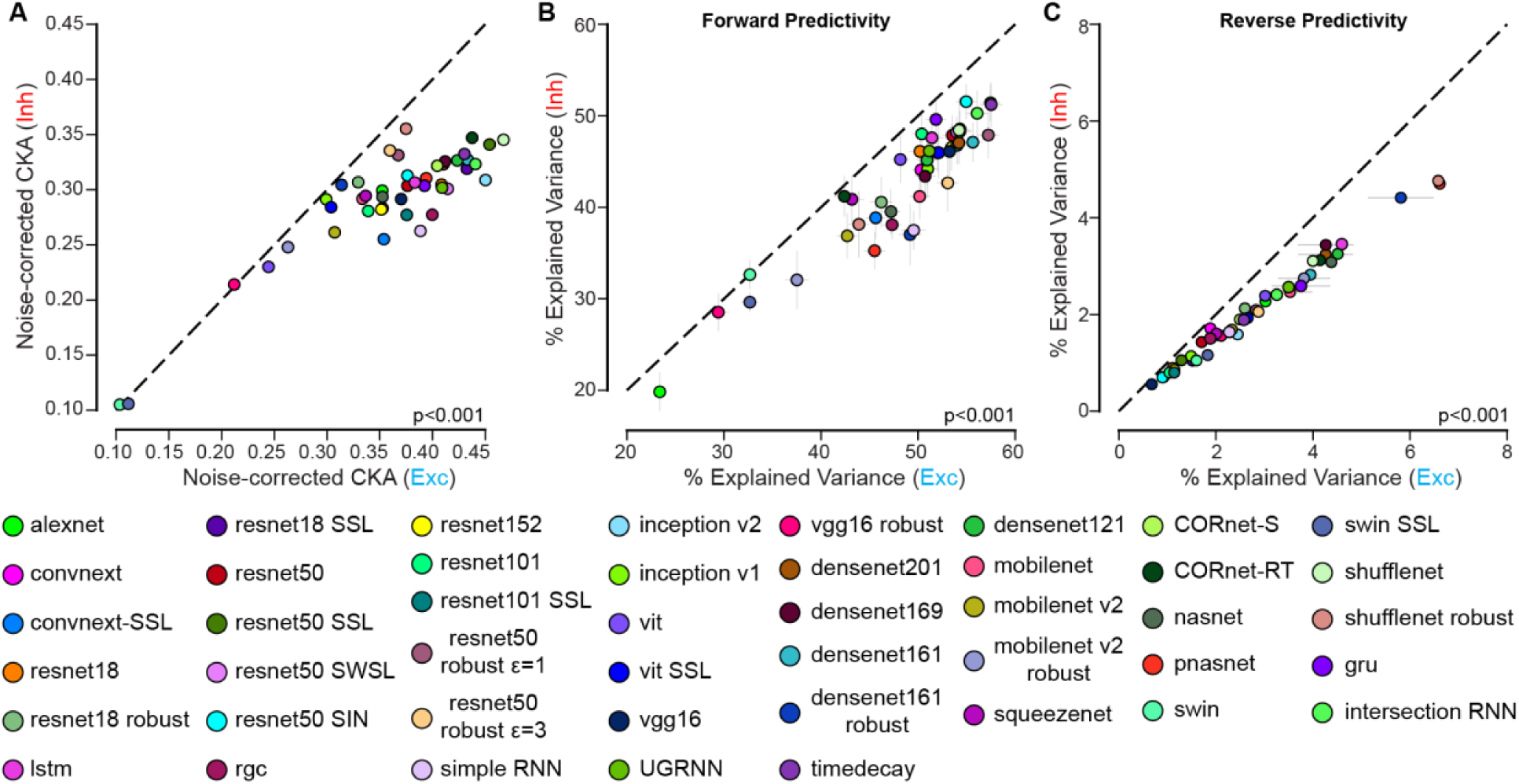
Model alignment with excitatory and inhibitory neural populations. **(A)** Noise-corrected CKA similarity between model representations and inhibitory (y-axis) versus excitatory (x-axis) neural responses. Each point represents one ANN model (color-coded by model). Models show higher alignment with excitatory populations than inhibitory populations (two-sided paired Wilcoxon test: *z=3*.*0, p < 0.001*). **(B)** Forward predictivity: percentage of explained variance when predicting neural responses from model features. Explained variance for inhibitory neurons (y-axis) is plotted against excitatory neurons (x-axis) for each model. Models explained more variance in excitatory than inhibitory responses (two-sided paired Wilcoxon test: *z=0.0, p < 0.001*). **(C)** Reverse predictivity: percentage of explained variance when predicting model features from neural responses. Explained variance for inhibitory neurons (y-axis) is plotted against excitatory neurons (x-axis). Reverse predictivity was consistently higher for excitatory populations across models (two-sided paired Wilcoxon test: *z=0.0, p < 0.001*).

**Figure S12.**
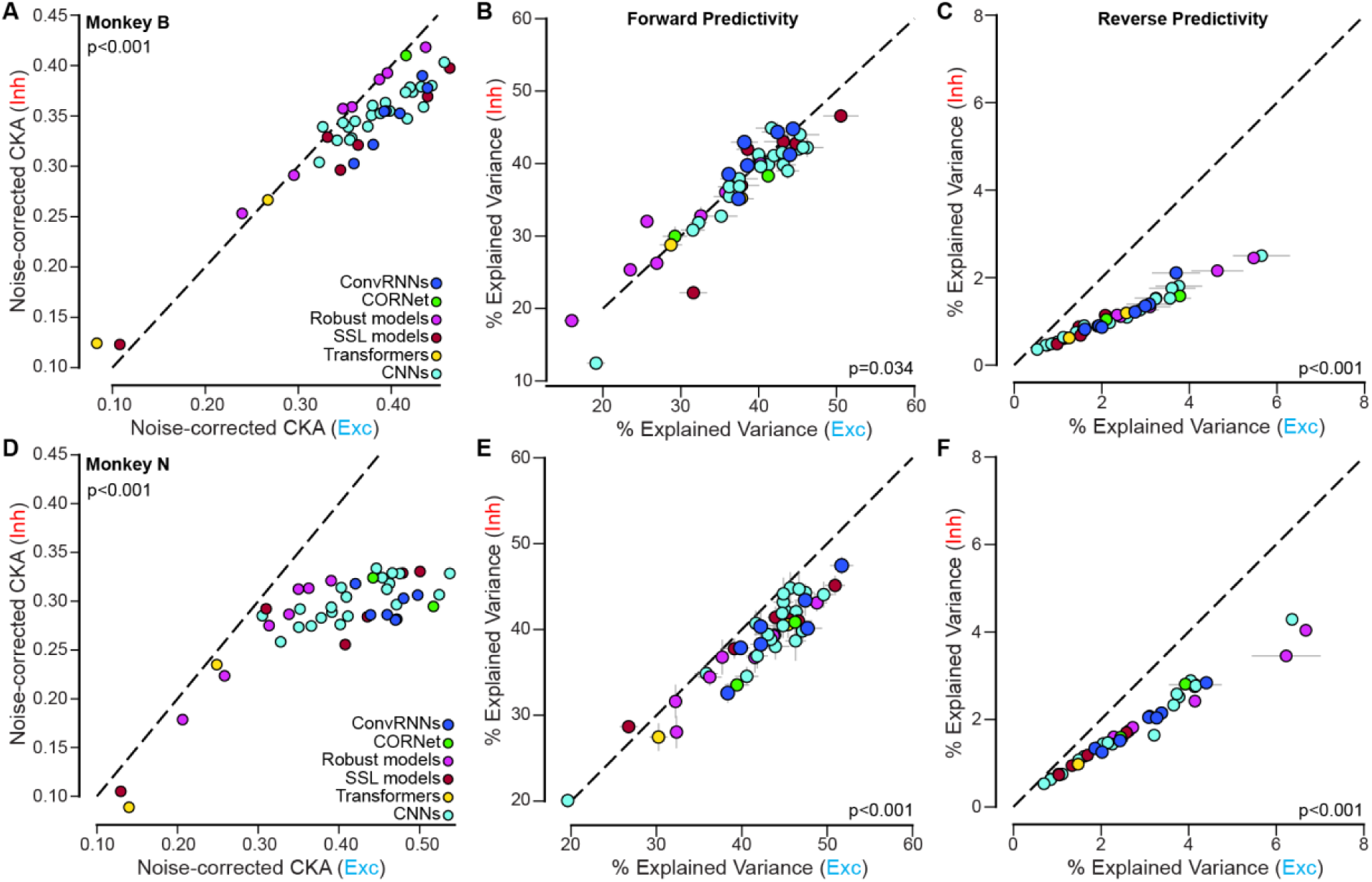
Model alignment with excitatory and inhibitory populations across monkeys. (**A**) Noise-corrected CKA similarity between model representations and inhibitory (y-axis) versus excitatory (x-axis) neural responses for Monkey B. Each point represents one ANN model (color-coded by architecture class). Models showed stronger alignment with excitatory than inhibitory populations (two-sided paired Wilcoxon test: *z=0.0, p < 0.001*). (**B**) Forward predictivity for Monkey B: percentage of explained variance when predicting neural responses from model features. Explained variance for inhibitory neurons (y-axis) is plotted against excitatory neurons (x-axis). Models explained more variance in excitatory than inhibitory responses (two-sided paired Wilcoxon test: *z=679.0, p = 0.034*). (**C**) Reverse predictivity for Monkey B: percentage of explained variance when predicting model features from neural responses. Explained variance for inhibitory neurons (y-axis) is plotted against excitatory neurons (x-axis). Reverse predictivity was consistently higher for excitatory populations (two-sided paired Wilcoxon test: *z=0.0, p < 0.001*). (**D–F**) Same analyses as in (**A–C**) for Monkey N. (**D**) Noise-corrected CKA similarity (two-sided paired Wilcoxon test: *z=0.0, p < 0.001*); (**E**) forward predictivity (two-sided paired Wilcoxon test: *z=1022.0, p < 0.001*); (**F**) reverse predictivity (two-sided paired Wilcoxon test: *z=0.0, p < 0.001*).

**Figure S13.**
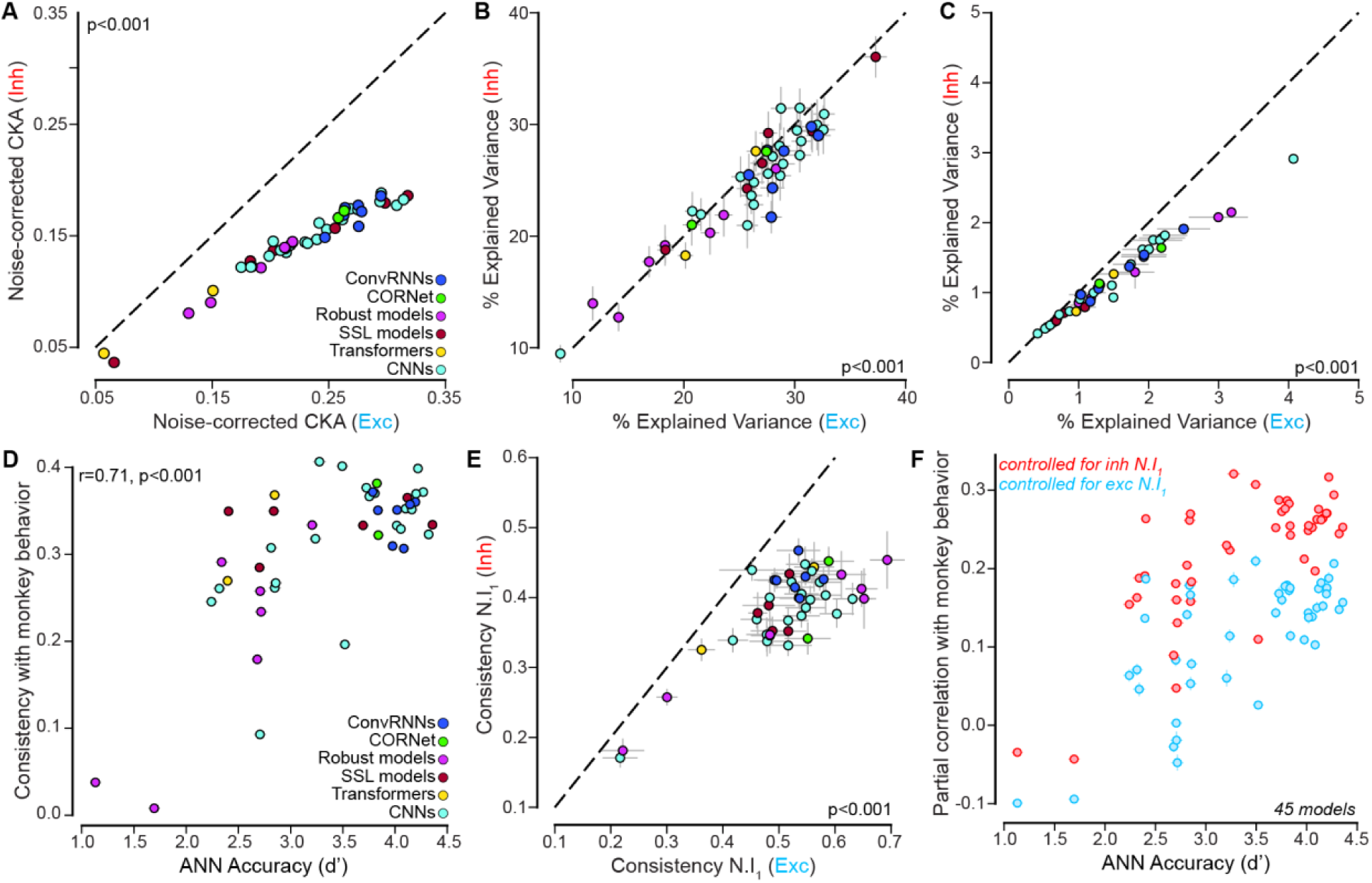
Relationship between model–neural alignment and behavioral consistency. **(A–C)** Same analyses as in Figure 7 and Figure S11 for the 180-280ms (time window leading the peak of the exclusive variance explained by Exc neurons). **(A)** Noise-corrected CKA similarity (two-sided paired Wilcoxon test: *z=0*.*0, p < 0.001*); **(E)** forward predictivity (two-sided paired Wilcoxon test: *z=202.0, p < 0.001*); **(F)** reverse predictivity (two-sided paired Wilcoxon test: *z=1.0, p < 0.001*). **(D-F)** Step-by-step variance partitioning analysis (Figure 8). (D) Relationship between overall ANN object recognition accuracy (d′) and consistency with monkey behavior. Each point represents one model (n=45 models). Behavioral consistency increased with model accuracy (noise-corrected Pearson correlation: *r(43) = 0*.*71, p < 0.001*). **(E)** Relationship between behavioral consistency derived from excitatory (x-axis) and inhibitory (y-axis) neural populations (N.I₁ metric). Excitatory-derived consistency exceeded inhibitory-derived consistency across models (two-sided paired Wilcoxon test: *z=0.0, p < 0.001*). **(F)** Partial correlation between model predictions and monkey behavior as a function of ANN accuracy. Red points show partial correlations controlling for inhibitory neural predictions; blue points show partial correlations controlling for excitatory neural predictions. Even after controlling for inhibitory predictions, models retained substantial correlation with behavior, whereas controlling for excitatory predictions markedly reduced behavioral correlation ((two-sided paired Wilcoxon test: *z=0.0, p < 0.001*). Each point represents one model (n = 45).

## Footnotes

† https://github.com/vital-kolab/reverse_pred/blob/main/lib/README.md

